# Designer antisense circRNA_GFP_ reduces GFP abundance in Arabidopsis protoplasts in a sequence-specific manner, independent of RNAi pathways

**DOI:** 10.1101/2023.11.20.567890

**Authors:** M Hossain, C Pfafenrot, S Nasfi, A Sede, J Imani, E Šečić, M Galli, P Schäfer, A Bindereif, M Heinlein, M Ladera-Carmona, KH Kogel

## Abstract

Circular RNAs (circRNAs) are single-stranded RNA molecules characterised by their covalently closed structure and are emerging as key regulators of cellular processes in mammals, including gene expression, protein function and immune responses. Recent evidence suggests that circRNAs also play significant roles in plants, influencing development, nutrition, biotic stress resistance, and abiotic stress tolerance. However, the potential of circRNAs to modulate target protein abundance in plants remains largely unexplored. In this study, we investigated the potential of designer circRNAs to modulate target protein abundance in plants using Arabidopsis as a model system. We demonstrate that treatment with a 50 nt circRNA_GFP_, containing a 30 nt GFP antisense sequence stretch, results in reduced GFP reporter target protein abundance in a dose- and sequence-dependent manner. Notably, a single-stranded open isoform of circRNA_GFP_ had little effect on protein abundance, indicating the importance of the closed circular structure. Additionally, circRNA_GFP_ also reduced GFP abundance in Arabidopsis mutants defective in RNA interference (RNAi), suggesting that circRNA activity is independent of the RNAi pathway. We also show that circRNA, unlike dsRNA, does not induce pattern-triggered immunity (PTI) in plants. Findings of this proof-of-principle study together are crucial first steps in understanding the potential of circRNAs as versatile tools for modulating gene expression and offer exciting prospects for their application in agronomy, particularly for enhancing crop traits through metabolic pathway manipulation.

**Highlights:** We demonstrate the potential of non-immunogenic circRNA as a tool for targeted gene regulation in plants, where circRNA acts in an isoform- and sequence-specific manner, paving the way for future agronomic applications.

## Introduction

Agricultural production is affected by a variety of biotic and abiotic stress factors, which will increase with higher temperatures and extreme weather conditions in the course of climate change (Pareek et al. 2020; IPPC Secretariat 2021). Further improvement or even maintenance of global yield levels will depend to a large extent on new scientific solutions and their rapid introduction into agronomic practice (Van Dijk et al. 2021). While there is a broad consensus in the scientific community and clear legal requirements in most countries that synthetic pesticides, including herbicides, should be used as little as possible (Deguine et al. 2021), the effectiveness of alternative crop protection measures in intensive production systems still needs to be developed, and their dependence on environmental factors is often poorly understood (Perez-Alvarez et al. 2019; Kremer et al. 2023; Galli et al. 2024).

RNA is key for the storage, transmission, and modification of genetic information. In higher organisms, RNA exists predominantly in the linear form as protein-coding mRNA and non-coding forms, such as ribosomal (r)RNAs, long non-coding RNA (lnc)RNAs, transfer (t)RNAs, and different types of small (s)RNA duplexes mostly of 21 to 24 base pairs. For the latter, their high significance for regulatory processes such as maintenance of genome stability and regulation of gene activity had only been found in 1998, when their function in RNA interference (RNAi) was discovered (Fire et al. 1998; Baulcombe 2004). As a way of communication between interacting organisms, RNA is also exchanged between animals or plants and their pathogens or parasites, a phenomenon known as cross-kingdom RNA interference (ckRNAi; LaMonte et al. 2012; Weiberg et al. 2013; Buck et al. 2014; Zhang et al. 2016; Shahid et al. 2018; for review see Cai et al. 2018; Hamby et al. 2025).

Consistent with the role of RNA in natural communication between plant hosts and microbial pathogens and pests, designer RNAs, such as engineered sRNA duplexes or longer double-stranded (ds)RNAs of up to several hundred nucleotides, can protect plants from biotic and abiotic stresses (for review see Koch and Kogel 2014; Cai et al. 2018; Niehl et al. 2018; Liu et al. 2020; Koch and Wassenegger 2021; Liu et al. 2024; Chen and Kim 2024). However, their instability and rapid degradation still hamper the agronomic use of these RNAs, especially if they are not protected by chemical formulations (Mitter et al. 2017; Demirer et al. 2019; Jain et al. 2022; Kogel 2025; Yong et al. 2025, Moorlach et al. 2025). Moreover, the risk of genetic cross-resistance to various sRNAs or dsRNAs all acting via RNAi in the target microbe or pest is a realistic scenario in which the RNAi pathway components and dsRNA uptake mechanisms are susceptible to counter-selection (Khajuria et al. 2018; OECD 2020; Wytinck et al. 2020; Mishra et al. 2021; Šečić and Kogel 2021; Choudhary et al. 2021; Luo et al. 2024; Mishra et al. 2024).

RNAs are also exchanged between plant hosts and their weed parasites (Westwood and Kim 2017). There is growing interest in exploring this potential use of RNA for weed control (Mai et al. 2021; Zabala-Pardo et al. 2022; Panozzo et al. 2025). However, RNA uptake and stability in plants have challenged the development of RNA herbicides (Dalakouras et al. 2016; Bennett et al. 2020; Liu et al. 2021; Yong et al. 2025), indicating the need for RNA with novel modes of action and molecular properties for their application in weed control.

In the present work, we have taken a first step to test the suitability of circular (circ)RNA for future agronomic applications. Unlike linear (lin)RNA molecules, circRNAs form a covalently closed loop, which confers resistance to exonucleases, making them more resistant to degradation (Nielsen et al. 2022; Liu et al. 2022). This circularization can be achieved through a process known as back-splicing, in which a downstream splice donor site joins with an upstream splice acceptor site, resulting in the formation of a closed loop. circRNAs can arise from exons (exonic circRNA), introns (intronic circRNA), and intergenic regions (Zhang et al. 2013; Jeck and Sharpless 2014). Knowledge about circRNAs has been generated mainly in animal systems, where they are involved in the regulation of gene expression at multiple levels, including their activity as microRNA (miRNA) sponges (binding to miRNAs and repressing their function), as protein scaffolds, or in sequestration and translocation of proteins, facilitation of interactions between proteins, or translation of proteins (Hansen et al. 2013; Memczak et al. 2013; Guo et al., 2014; Yang et al. 2022). As a result, circRNAs modulate various physiological processes such as cell differentiation, development, and cellular immune responses, and play a role in numerous diseases, including cancer and neurological disorders, with their therapeutic potential widely recognized (He et al. 2021; Pisignano et al. 2023; Liu et al. 2022; Guo et al. 2025). circRNAs also have been detected in plants, where they accumulate in response to biotic and abiotic stress (Zhang and Dai 2022; He et al. 2025). A comparison of 6,519 circRNAs from rice (*Oryza sativa*) with those from 46 other species revealed a high degree of conservation within the Oryza genus (46%), and as much as 8.5% were also found in dicotyledonous plants, indicating some conservation of circRNAs in plants (Chu et al. 2022). An endogenous antisense circRNA was reported to regulate the expression of the small subunit of RuBisCO in *Arabidopsis thaliana* (Zhang et al. 2021). Interestingly, Arabidopsis circRNAs have also been detected in leaf intercellular washing fluids (IWF), showing that they can be secreted to the plant apoplast where they potentially get in contact with plant attacking microbes (Zand Karimi et al. 2022). Notably, apoplastic circRNAs are highly enriched in the posttranscriptional modification N6-methyladenine (m6A), which is known to efficiently initiate circRNA translation (Yang et al. 2017).

Here we explore the potential of exogenously applied designer circRNAs to target an endogenous Green Fluorescence Protein (GFP) reporter protein in Arabidopsis. *GFP*-expressing cells treated with the GFP-specific circRNA_GFP_, in contrast to its corresponding linear single-strand form linRNA_GFP_ or a circRNA that does not contain GFP-specific target sequences (circRNA_CTR1_), showed reduced GFP protein abundance in a sequence- and circRNA-isoform-specific manner. Moreover, using RNAi mutants compromised in DICER-LIKE (DCL) and ARGONAUTE (AGO) activities, we demonstrate that the circRNA-mediated activity on reporter protein abundance is independent of the canonical RNAi pathways.

## Results

### Design of *GFP*-antisense circRNA

In a previous study, Pfafenrot and co-workers (2021) showed in the mammalian system that antisense-circRNAs can be designed to efficiently interfere with translation of a protein-coding gene. To develop a new tool for targeting gene expression with exogenous RNA, we synthesized circRNA targeting the ORF of a *GFP* reporter gene (Fig. S1A). The exact position of the target sequence was selected based on the secondary structure model of the ORF (Fig. S1B). The selection of this region was confirmed by measuring mRNA accessibility using the RNAup webtool (Fig. S1C). Based on this information, we designed a 50 nucleotide (nt) long antisense circRNA (circRNA_GFP_) that contained a central anti-*GFP* sequence of 30 nt with perfect complementarity. In addition, two different non-specific circRNAs were synthesized, which contained a randomized 25 nt or 46 nt sequence with a common 20 nt backbone, forming 45 nt circRNA_CTR1_ and 66 nt circRNA_CTR2_, respectively. Secondary structure models of all circRNAs are shown in Fig. S1D (for sequences, see Table S1).

### circRNA_GFP_ reduces the GFP abundance in *GFP*-expressing protoplasts in a sequence-specific manner

To evaluate the antisense activity of the designed circRNAs, mesophyll protoplasts isolated from Arabidopsis leaves were cotransfected with 4 μg of circRNA_GFP_ or the non-specific circRNA_CTR1_ and 20 μg of plasmid pGY1-35S::GFP:RFP (Fig. S2A). After 18 h of incubation (hpt) in the dark, the transfected protoplasts were analysed by measuring the ratio of GFP fluorescence to RFP fluorescence using ImageJ. Notably, we found that the GFP fluorescence was significantly reduced only in the circRNA_GFP_-treated sample as compared to the circRNA_CTR1_ or the untreated controls (Fig. 1A, B). To further substantiate this finding, we used an alternative GFP-expressing plasmid to transfect protoplasts. Similarly and consistent with our expectation, in protoplasts transfected with pGY1-35S::GFP (Fig. S2B), GFP fluorescence was also reduced upon treatment with circRNA_GFP_, but not in samples treated with circRNA_CTR1_ or untreated controls, when normalized to red chlorophyll autofluorescence (Fig. S3A, B). This finding indicated that circRNA_GFP_ exerted an inhibitory effect on GFP abundance in a sequence-specific manner.

**Figure 1.**
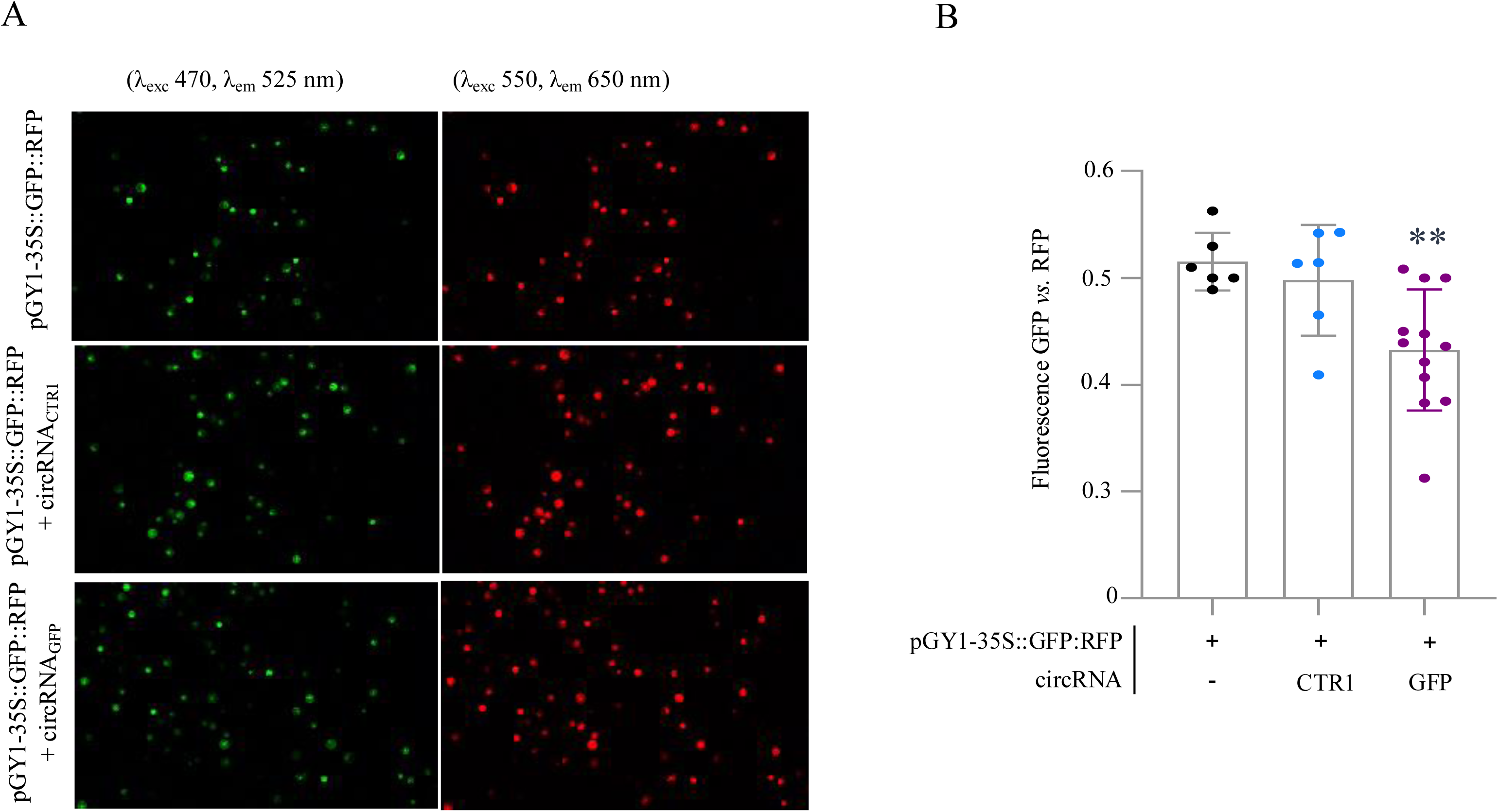
Microscopic imaging of GFP and RFP fluorescence in Arabidopsis protoplasts. Protoplasts were transfected with 20 µg of plasmid pGY1-35S::GFP:RFP and 4 µg of *GFP* antisense circRNA_GFP_, or 4 µg of non-targeting circRNA_CTR1_. (A). After 18 hpt, protoplasts were examined under the microscope using two distinct filters to GFP fluorescence (λexc 470, λem 525 nm) and RFP fluorescence (λexc 550, λem 650 nm). Fluorescence intensity was quantified based on images by using ImageJ 1.54p software. (B). The ratio between green pixels (GFP fluorescence) and red pixels (RFP fluorescence) as calculated with ImageJ is represented in the graph. The bar represents the measurements of ≥6 individual pictures taken at various positions. Statistical analysis was performed using one-way ANOVA, followed by Dunnett’s multiple comparison test, where ‘**’ denotes p ≤ 0.01 significance to the protoplast transfected without circRNA (control). Bars show standard deviation (SD).

### The impact of various doses of circRNA_GFP_ on GFP abundance

To further confirm target specificity of the designed circRNA, Arabidopsis protoplasts were cotransfected with 20 µg pGY1-35S::GFP and increasing amounts of circRNA_GFP_ and circRNA_CTR1_. ImageJ analyses of the GFP fluorescence after 18 hpt indicated that the effect of circRNA_GFP_ was slightly concentration dependent and remained circRNA-sequence-specific over a concentration range up to 8 µg (Fig. 2A; Fig. S4A,B). Next, we quantified the effect of circRNA_GFP_ on GFP abundance in protoplasts by immunoblot analyses. For this purpose, protoplasts were isolated from stable, transgenic *GFP*-expressing Arabidopsis plants and subsequently treated with increasing concentrations of circRNA_GFP_ and circRNA_CTR1_. At 18 hpt, total protoplast proteins were extracted and separated by gel electrophoresis. GFP protein abundance was visualised after blotting with an anti-GFP antibody and an anti-actin antibody was used for protein normalisation. Consistent with the fluorescence analyses, we found that the amount of GFP was reduced in protoplasts treated with increasing concentrations of circRNA_GFP_, as compared to protoplasts treated with circRNA_CTR1_ (Fig. 2B).

**Figure 2.**
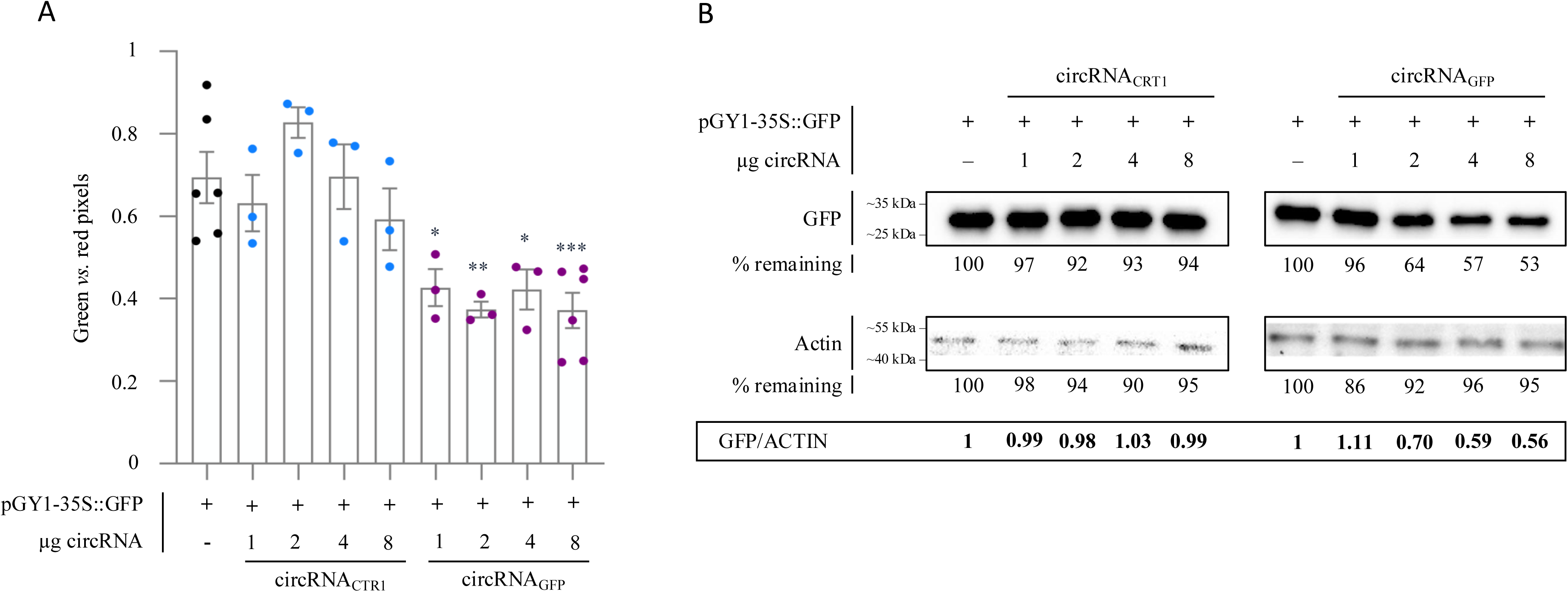
Dose dependence of GFP fluorescence in Arabidopsis protoplasts after treatment with increasing concentrations of circRNAs. Protoplasts were cotransfected with 20 µg of plasmid pGY1-35S::GFP and the indicated amounts of circRNA_GFP_ or circRNA_CTR1_. (A) After 18 hpt, protoplasts were inspected under the fluorescence microscope using two different filters to calculate the ratio in fluorescence levels between the GFP fluorescent protoplasts (λ_exc_ 470, λ_em_ 525 nm) and total protoplasts (red autofluorescence from the chlorophyll, λ_exc_ 480, λ_em_ 510 nm). Fluorescence was measured based on pictures by using ImageJ 1.54p software. Bars represent the average of the measurements of at least 3 pictures taken at different spots with standard error of the mean (SEM). Statistical analysis was performed using one-way ANOVA, where stars denote significance to the protoplast transfected without circRNA (control). (Dunnett’s test). ‘*’ p ≤ 0.05, ‘**’ p ≤ 0.01, ‘***’ p ≤ 0.001. (B) Immunoblot analysis of proteins extracted from protoplasts treated with increasing concentrations of circRNA_GFP_ or circRNA_CTR1_. The values below the GFP band indicate the remaining amount of GFP protein in percent as detected using an anti-GFP antibody. An anti-Actin antibody was used to visualize Actin as an internal loading control. The ratio of GFP protein accumulation is indicated below the bands expressed as ratio between GFP vs. Actin signals).

### The impact of circRNA_GFP_ on GFP abundance is isoform-specific

Next, we comparatively examined the effect circRNA and its single-stranded, non-circularised linear antisense form (linRNA_GFP_) on GFP abundance, where linRNA_GFP_ consisted of the same nt sequence as circRNA_GFP_. For this purpose, Arabidopsis wild-type protoplasts were treated with the plasmid pGY1-35S::GFP and 4 µg of either circular (circRNA_GFP_) or linear (linRNA_GFP_) configurations of the GFP antisense RNA, and incubated for 10 h, 18 h, and 32 h. circRNA_GFP_-mediated inhibition of GFP abundance was already detectable at 10 h after protoplasts treatment, and this effect persisted until 32 h (Fig. 3). In contrast, linRNA_GFP_ showed a transient inhibitory effect on protein abundance after 10 h, which disappeared over time. Overall, our analyses revealed an isoform-specific effect of circRNA_GFP_ on GFP abundance. Our data suggest that circRNA is more effective than its corresponding single-stranded linear RNA in antisense targeting of plant gene expression.

**Figure 3.**
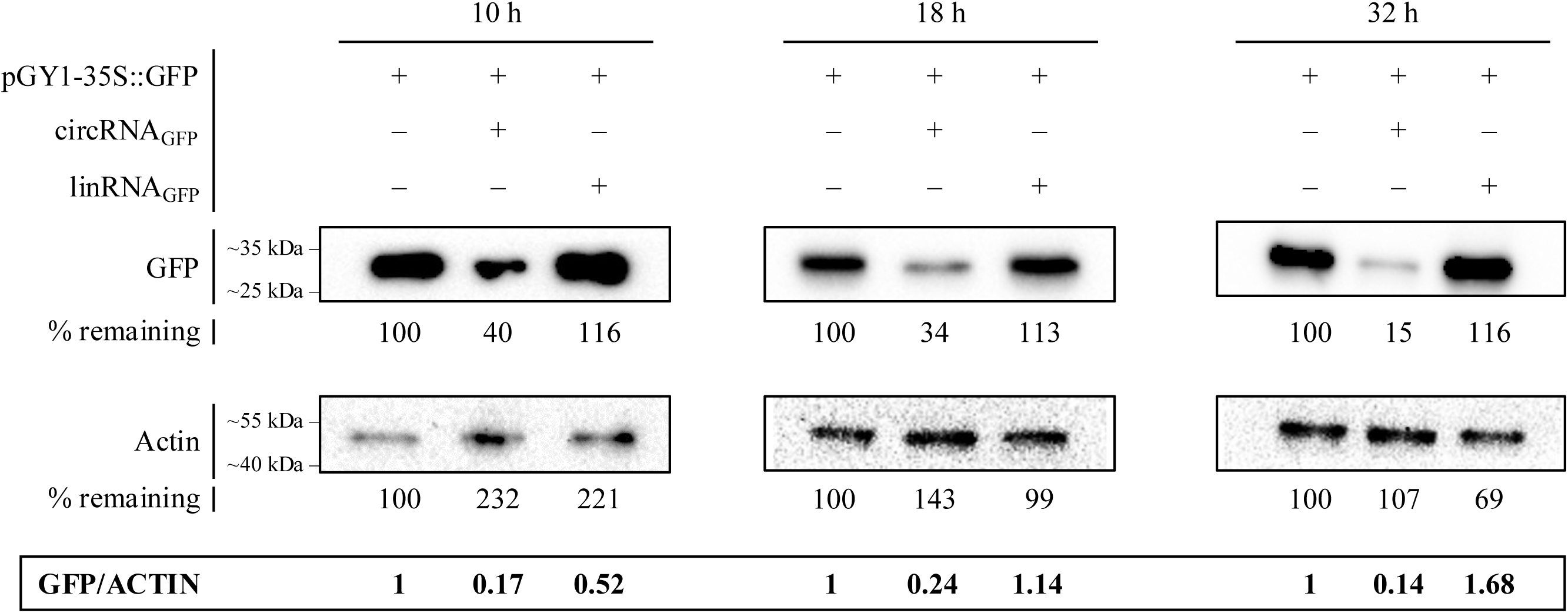
Immunoblot analysis of GFP abundance in Arabidopsis protoplasts after treatment with the *GFP* antisense circRNA_GFP_ or its corresponding single-stranded, linear form linRNA_GFP_. Protoplasts were transfected with pGY1-35S::GFP and 4 µg circRN_GFP_ or linRNA_GFP_ and analysed for GFP abundance after 10 h, 18 h and 32 h after transfection using an anti-GFP antibody. An anti-actin antibody was used to visualize actin as a loading control. GFP abundance is indicated below the band as percentage of GFP in protoplasts not treated with RNA (control). The ratio of GFP protein abundance is indicated below the bands expressed as ratio between GFP vs. actin signals).

### circRNA_GFP_ affects GFP abundance independently of functional DCLs and AGOs

To obtain further information on the mode of action of sequence-specific designer circRNA on target protein abundance, we investigated whether the observed effect of circRNA_GFP_ was lost in Arabidopsis mutants impaired in RNAi. Accordingly, protoplasts isolated from Arabidopsis DCL and AGO mutants were cotransfected with 20 μg plasmid pGY1-35S::GFP and 4 μg of the respective circRNA. Like wild-type protoplasts, *dcl1-11* and *ago1-27* protoplasts showed reduced GFP fluorescence in response to circRNA_GFP_ (Fig. 4A,B; Fig. S5A,B), suggesting that disruption of DCL1 and AGO1 activities had no effect on circRNA_GFP_-mediated reduction in GFP protein abundance. Consistent with this finding, immunoblot analyses further confirmed that disruption of the RNAi pathway did not affect the circRNA_GFP_ effect. We found reduced GFP abundance in circRNA_GFP_-treated *dcl1-11* (57%), *ago1-27* (49%) and wild-type (78%) protoplasts as compared to untreated protoplasts, whereas circRNA_CTR1_ did not affect GFP abundance in either wild-type or mutant protoplasts (Fig. 4C). Immunoblot analyses of GFP abundance in additional RNAi mutants, including the DCL triple mutant *dcl2,3,4* and the two AGO mutants *ago2-1* and *ago4-2*, further substantiated that circRNA_GFP_ retained its effect on GFP abundance when the RNAi pathway was compromised (Fig. S6).

**Figure 4.**
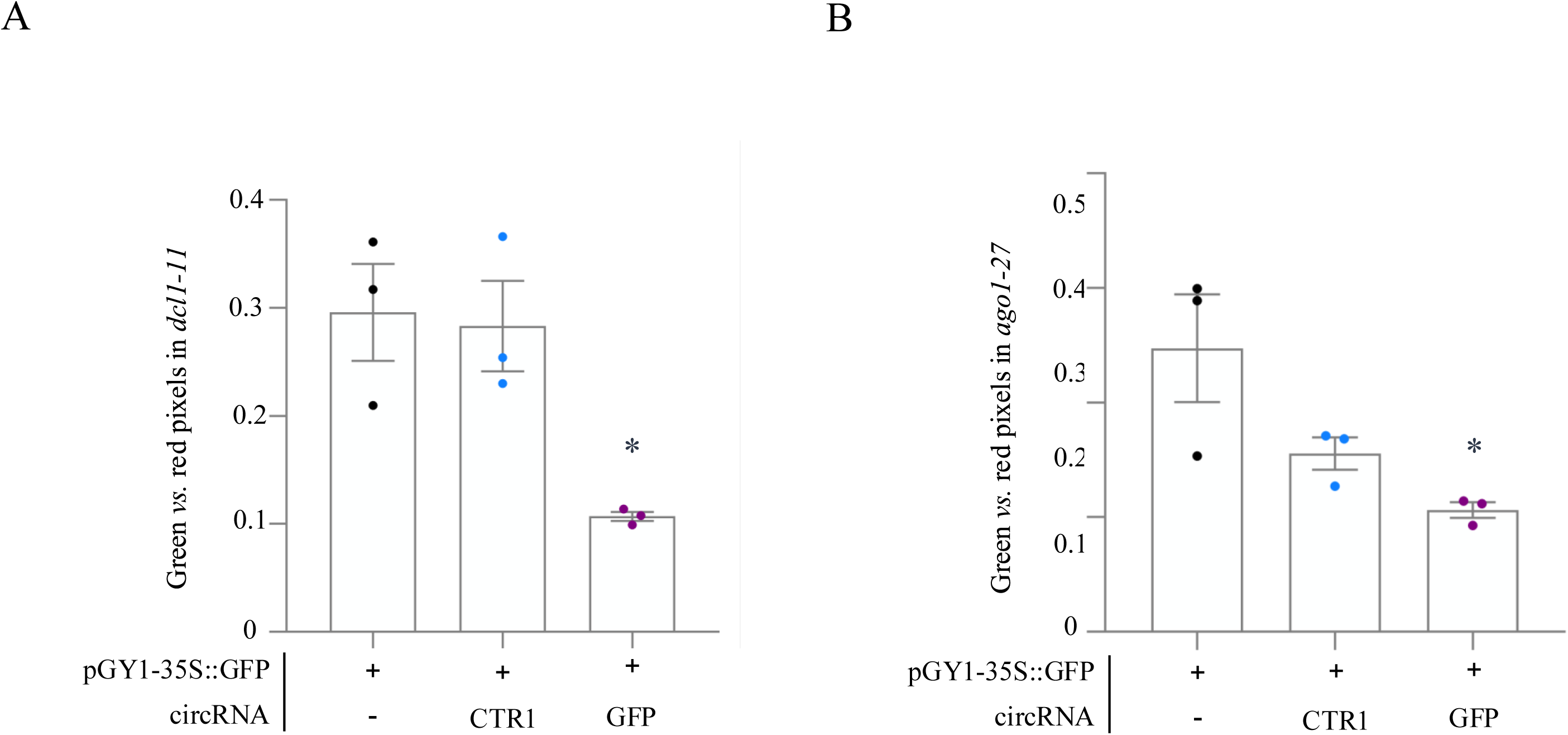
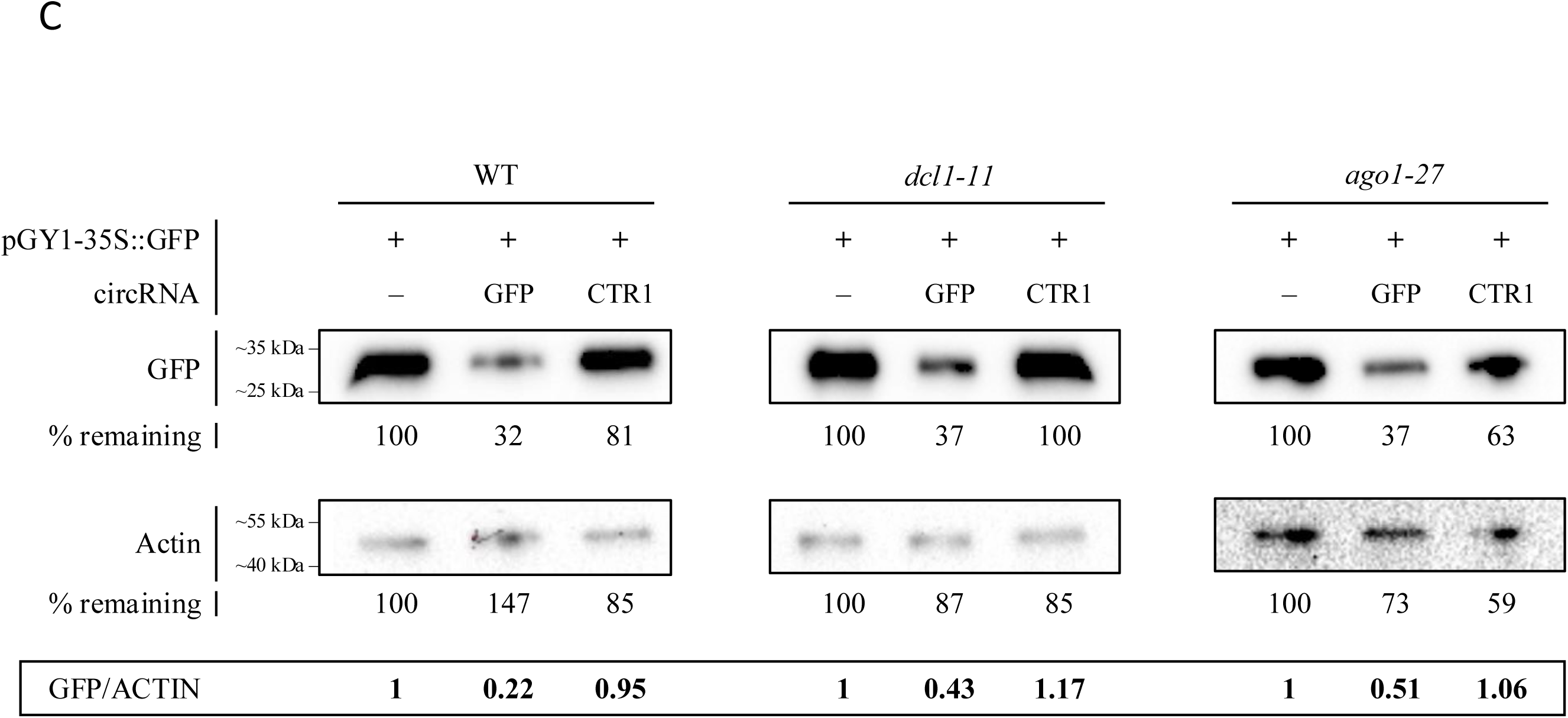
GFP abundance in Arabidopsis protoplasts of RNAi mutants *dcl1-11* and *ago1-27* after circRNA treatment. (A, B) Imaging of GFP fluorescence in *dcl1-11* (A) and *ago1-27* (B) at 18 hpt. Protoplasts were transfected with 20 µg of plasmid pGY1-35S::GFP and 4 µg of *GFP* antisense circRNA_GFP_ or non-targeting circRNA_CTR1_. Depicted is the ratio between green pixels (GFP fluorescence) and red pixels (chlorophyll fluorescence) as calculated with ImageJ. Bars represent the average of the measurements of at least 3 pictures taken at different spots with standard error of the mean (SEM). Statistical analysis was performed with one-way ANOVA, where * denotes p≤0.05 significance vs. control protoplasts (Dunnett’s test). (C). Immunoblot analysis of the GFP abundance in protoplasts from wild-type and RNAi mutants *dcl1-11* and *ago1-27* upon treatment with circRNA_GFP_ or circRNA_CTR1_. Protoplasts were harvested at 18 hpt, and equal amounts of protein were analyzed by immunoblotting, using an anti-GFP antibody and an anti-Actin antibody as a loading control. The ratio of GFP abundance is indicated below the bands expressed as ratio between GFP vs. Actin signals.

### circRNA_GFP_ has no impact on *GFP* transcript abundance

Next, we analyzed the effect of circRNA_GFP_ on the level of the *GFP* transcript in transgenic Arabidopsis protoplasts. Based on our previous work (Pfafenrot et al. 2021) we hypothesized that target mRNA levels would not be reduced upon circRNA treatment. To this end, *GFP*-expressing protoplasts were transfected with 20 µg pGY1-35S::GFP:RFP alone or together with either 4 µg circRNA_GFP_ or circRNA_CTR1_/circRNA_CTR2_ followed by measurements of *GFP* transcript levels at 18 hpt. RT-qPCR analyses showed that none of the circRNAs reduced *GFP* transcript levels significantly in the wild-type (Fig. 5), and in all the mutants comprising *dcl1-11*, *ago1-27*, *dcl2,3,4*, *ago2-1* and *ago4-1*. (Fig. S7A-E). These findings showed that circRNA_GFP_ inhibited protein abundance in a sequence- and isoform-specific manner without affecting *GFP* transcript levels, through a process that was independent of canonical RNAi pathways.

**Figure 5.**
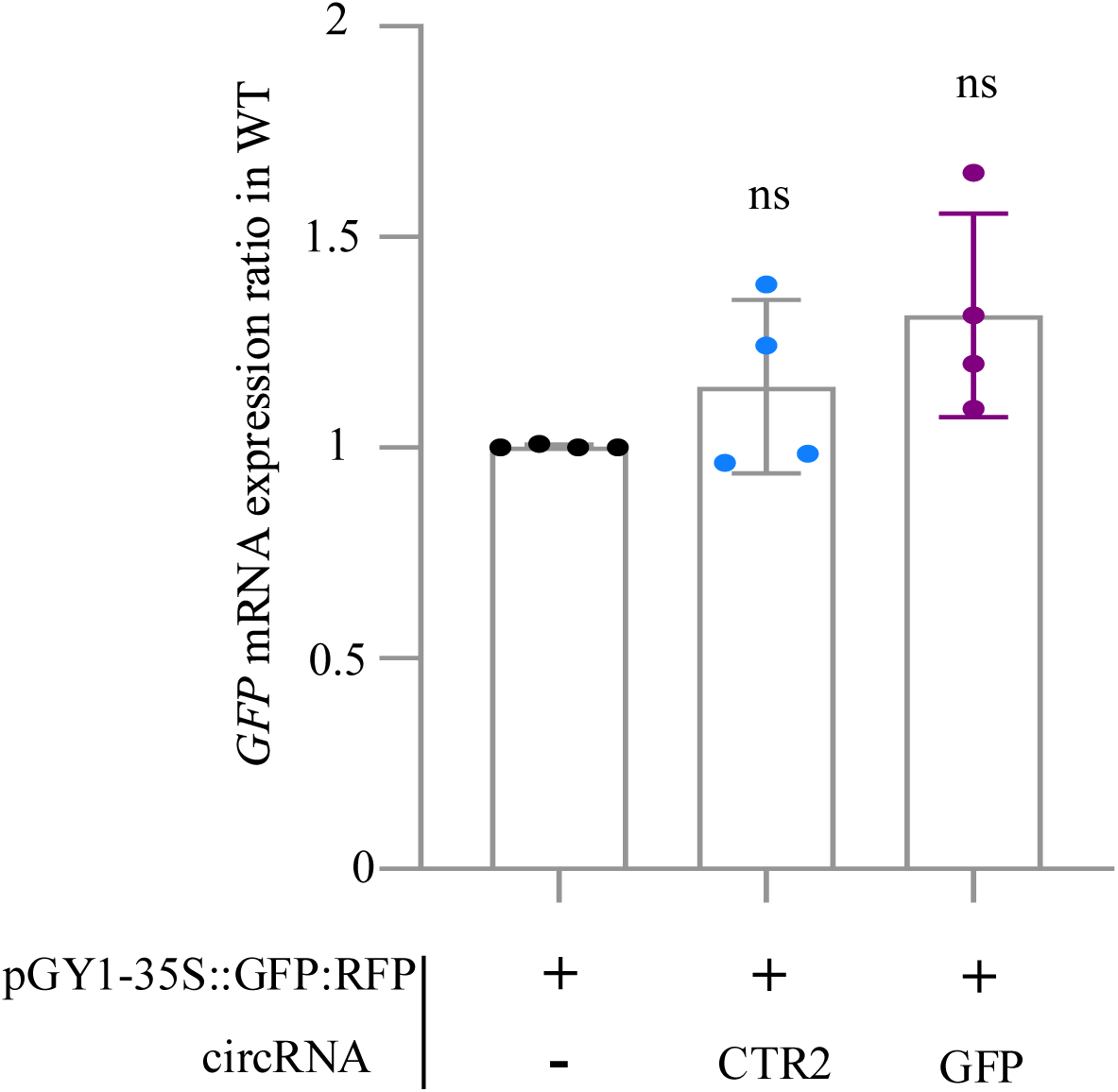
RT-qPCR analysis of the amount of *GFP* transcripts in Arabidopsis wild-type protoplasts upon treatment with circRNA. Protoplasts were transfected with 20 µg of pGY1-35S::GFP:RFP plasmid and 4 µg of circRNA_GFP_ or non-targeting circRNA_CTR2,_ respectively. Relative *GFP* expression was measured after 18 hpt. Values were normalized to RFP and the housekeeping genes *Ubiquitin* and *EF1-*α. Bars represent an average of three independent biological experiments pooled together with standard deviation (SD). No statistically significant differences between treatments and genotypes were detected (Kruskal-Wallis test).

### circRNA does not induce typical PTI responses in leaves

dsRNA activates pattern-triggered immunity (PTI) in plants leading to various responses, including callose deposition at plasmodesmata and MAP kinase activation (Niehl et al. 2016; Huang et al. 2023). We wondered whether similar to dsRNA also circRNA triggers a PTI response. To this end, equal molar amounts of circRNAs (circRNA_CTR1_, circRNA_CTR2_), their corresponding linear forms linRNA_CTR1_ and linRNA_CTR2_, or the synthetic dsRNA analog poly(I:C) (as positive control) were vacuum-infiltrated into Arabidopsis leaf disks together with aniline blue to stain callose. Fluorescence microscopy revealed that poly(I:C) treatment induced a strong aniline blue fluorescence at the plasmodesmata, as compared to the other treatments (Fig. 6A, B). Quantification of fluorescence showed that poly(I:C) triggered the highest mean callose intensity. Equal molar amounts (corresponding to 50 ng µL^−1^) of the linear form of RNA_CTR1_ (linRNA_CTR1_) also triggered callose deposition, although to a lesser extent. Surprisingly, neither of the circRNA molecules induced an increased callose intensity, even when the circRNA concentration was increased by 5 times to 250 ng µL^−1^ (Fig. S8). In line with this, immunoblot analyses to detect mitogen-activated protein kinase (MAPK) phosphorylation in *Arabidopsis thaliana* leaves using anti-phospho-p44/42 ERK antibodies consistently revealed a strong induction of MAPK phosphorylation in response to 1 µM flg22, but not upon treatment with ∼3 µM linRNA_CTR1_ or circRNA_CTR1_ (Fig. 6C). Finally, to assess whether circRNAs trigger a reactive oxygen species (ROS) response, *Nicotiana benthamiana* leaf discs were treated with 1 µM flg22, ∼1 µM poly(I:C), or ∼1 µM linRNA_CTR2_ or circRNA_CTR2_. In line with expectations, flg22 elicited a robust ROS burst, whereas poly(I:C), linRNA_CTR2_, and circRNA_CTR2_ failed to induce ROS accumulation (Fig. 6D). These results support the possibility that, unlike dsRNA, circRNAs may be able to evade receptor recognition in plants.

**Figure 6.**
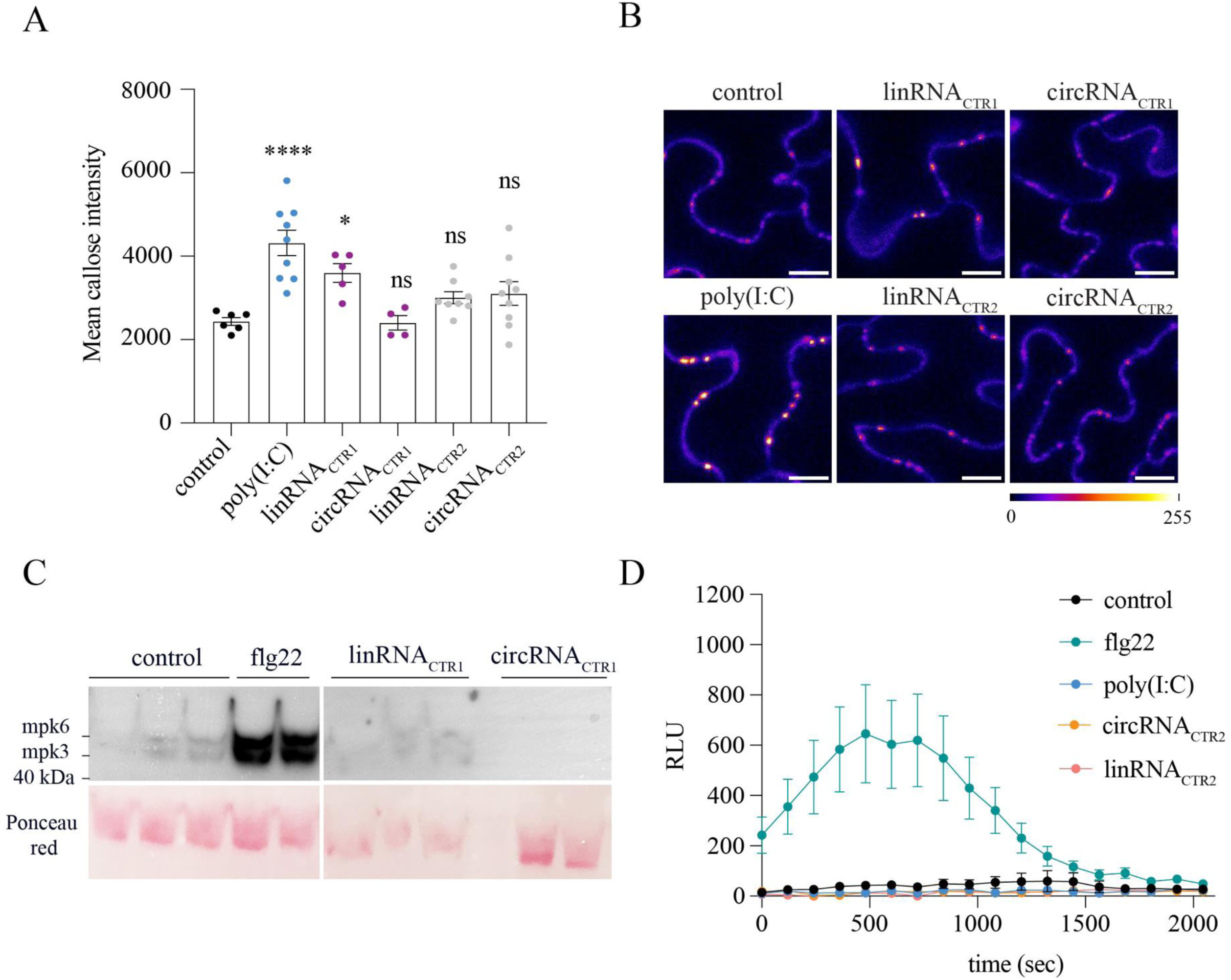
Analysis of immune innate responses after treatment with various unspecific RNAs. **(A)** Mean values of callose intensity levels at PD in individual images taken of epidermal cells of leaf disks treated with a 0.1% aniline blue solution containing water or 500 ng/µL of poly(I:C), or 50 ng/µL of either linRNA or circRNA. Error bars show the standard error of the mean. Parametric mean value variances were tested by one-way ANOVA followed by Dunnett’s multiple comparisons test. ****, p< 0.0001; *, p<0.05; ns, non-significant. **(B)** Callose accumulation at plasmodesmata (PD) in Arabidopsis leaves upon staining with aniline blue. Samples were treated with 0.1% aniline blue solution together either with water (negative control) or 500 ng/µL of poly(I:C) (positive control), or with 50 ng/µL of either linRNA_CTR1_, circRNA_CTR1_, linRNA_CTR2_, or circRNA_CTR2_. For a better visualization, color was assigned to the 8-bit images by applying the “fire” look-up table using the Image J software. Scale bar, 10 µm. **(C)** Immunoblots for the detection of mitogen-activated protein kinase (MAPK) phosphorylation in Arabidopsis leaves probed with antibodies against phosphor-p44/42 ERK. Leaf disks were vacuum infiltrated with either water (negative control), 1 µM flg22 or ∼3 µM of each linRNA_CTR1_ or circRNA_CTR1_. As control for equal loading, Ponceau Red staining was used. **(D)** ROS production assay in *N. benthamiana* leaf discs after incubation with either water (negative control), 1 µM flg22, ∼1 µM poly(I:C) or ∼1 µM of linRNA_CTR2_ or circRNA_CTR2_. Mean values and the standard error of the mean (SEM) are shown from nine independent replicates per treatment. RLU, relative luminescence units.

## Discussion

The use of exogenous RNA to directly influence gene expression in crops is understudied despite its potential applicability as, for example, selective herbicides or as antimicrobial agents targeting plant susceptibility genes (Zabala□Pardo et al., 2022, Mai et al., 2021; van Schie and Takken, 2014). Current RNA strategies based on dsRNA and sRNA, are challenged by low stability upon leaf application under field conditions (Mitter et al. 2017; Bachman et al. 2020; Kogel 2025; Yong et al. 2025). This limitation prompted us to conduct a baseline study to investigate whether more stable circular RNAs have properties that could make them a potentially effective additional alternative in crop protection in the future. Our findings demonstrate that exogenously applied circRNA containing an antisense sequence to a *GFP* reporter gene (circRNA_GFP_) significantly reduces GFP protein abundance in Arabidopsis cells. This effect is *i.* topology-dependent as the circular configuration is crucial, *ii.* sequence-specific as non-targeting control circRNA was inactive, *iii.* RNAi-independent, as responses to circRNA were similar in wild-type and RNAi-deficient mutants, and *iv.* accordingly different from the mode of action of dsRNA. Overall, these results suggest that circRNA application for manipulation of plant gene and metabolism is feasible, justifying further fundamental research for future practical application.

circRNA is generally more stable than dsRNA due to its intrinsic physical properties: *i.* circRNA has a covalently closed-loop structure, which prevents degradation by exonucleases – unlike dsRNA, which has free 5’ and 3’ ends that are susceptible to degradation; *ii.* circRNA is resistant to most RNA-degrading enzymes that typically target linear RNA molecules, while dsRNA can be degraded by endonucleases such as RNase III; *iii.* circRNA has a high secondary structure stability and can maintain its integrity under harsher conditions, while the stability of dsRNA is affected by environmental factors such as temperature and pH (Liu and Chen 2022; Nielsen et al. 2022; Ren et al. 2022; Liu et al. 2022; Moorlach et al. 2025). Future studies are needed to determine whether the physical stability of circRNA makes it more suitable for use in crop protection, considering both efficacy and environmental safety implications.

dsRNA can trigger immune responses in animals, plants and fungi (de Reuver and Maelfait 2024; Niehl et al. 2016; Zheng et al. 2025). circRNA, on the other hand, is known to be less immunogenic in mammals (Wesselhoeft et al. 2019). In line with this, we show that circRNA does not induce callose deposition, ROS accumulation or MAP kinase activity, three hallmarks of PTI responses in plants, whereas the dsRNA analogue poly(I:C) induced robust callose deposition at plasmodesmata and MAP kinase (see Figs. 6 and S8). These results also confirm earlier reports that dsRNA does not trigger ROS in plants (Niehl et al. 2016; Huang et al. 2023). These observations, in line with reports of circRNAs not triggering antiviral responses in mammalian cells (Breuer et al. 2022), highlight the specificity of circRNAs and suggests they are less likely to provoke off-target immune reactions. This biological property is also an interesting aspect when considering the potential of circRNA as a crop protection agent.

We used the Arabidopsis protoplast system as the first proxy for evaluating the effect of circRNAs on target proteins in plants. To make the protoplast experiments robust, we used two different reporter constructs for transfection in independent experiments. Both constructs pGY1-35S::GFP:RFP and pGY1-35S::GFP were expressed in the protoplasts and targeting the GFP reporter by circRNA_GFP_ resulted in a reduction of GFP fluorescence in both cases. Consistent with this observation, immunoblot-based assays confirmed that the exogenous application of circRNA_GFP_, but not circRNA_CTR1_, reduced the abundance of GFP protein.

The observation that linRNA_GFP_, in contrast to circRNA_GFP_, only transiently downregulated GFP abundance (see Fig. 3) is consistent with the observation by our earlier work (Pfafenrot et al. 2021), where the inhibitory potency of circular forms of antisense sequences consistently surpassed their linear version. Therefore, the lasting inhibitory effect of circRNAs is likely due to the high metabolic stability over linear forms, being more resistant to the attack of exonucleases (Pfafenrot et al. 2021).

We also show here that the sequence-specific activity of circRNA occurs at the translation level, as the mRNA-GFP transcript level is not reduced, while the GFP protein level decreases significantly. That circRNA acts on the translational level rather than affecting target transcript abundance is also consistent with Pfafenrot et al. (2021), who showed that various designer circRNAs targeted viral RNA in infected mammalian cells in a sequence-specific manner. The possibility that endogenous circRNA act on RNA transcripts in mammalian systems has been recently discussed (Wang et al. 2024). Based on these recent reports along with our findings we suggest a mode of circRNA action in plant cells where the circRNA binds in a sequence-specific manner on target mRNAs and therefore inhibits or delays translation. The slightly (albeit insignificantly) higher transcript abundance of the target *GFP* mRNA in circRNA_GFP_-treated samples could indicate that the transcript may be stabilized by the binding.

Protoplasts have no cell walls and PEG facilitates the penetration of plasmids and other nucleic acids into eukaryotic cells (Wu et al. 2009). By using protoplasts, we were able to circumvent the problem of topical application of RNA on plant leaves, which is still a major technical challenge due to the numerous barriers that need to be overcome, including the plant cuticle and cell wall (Bennett et al. 2020; Kogel 2025). The issue of uptake of circRNA by spray application in the field is one that requires further significant scientific input. While there is no robust data on circRNA uptake through plant leaves, it is possible that future applications may require formulations or physical means, as has been shown for dsRNAs and small RNAs (Dalakouras et al. 2016; Mitter et al. 2017; Demirer et al. 2019; Yong et al. 2025).

In summary, this study demonstrates that exogenously applied designer circRNAs can regulate protein expression in plants through a sequence-specific activity and an RNAi-independent mode of action. These results pave the way for future studies aimed at using circRNAs to develop new RNA-based herbicides and antimicrobials.

## Experimental Procedures

### Plant material and isolation of Arabidopsis protoplasts

*Arabidopsis thaliana* and *Nicotiana benthamiana* plants were grown from seeds in soil (LAT-Terra Standard Topferde Struktur 1b, Hawita, Vechta, Germany) complemented with 2,5 g/L fertilizer (Osmocote 12-7-19 +TE) at 22°C/18°C under 12 h/12 h or 16 h/8 h, light:dark cycles, respectively. *A. thaliana* (Col-0) wild-type (WT), the mutants *ago1-27*, *ago2-1*, *ago4-1*, *dcl1-11* and the triple mutant *dcl2,3,4* were obtained from NASC (https://arabidopsis.info/). All mutants were verified by genotyping. *A. thaliana* plants constitutively expressing *GFP* were published in Harvey *et al*. (2020). Mesophyll protoplasts were produced using the tape-Arabidopsis-sandwich method (Wu *et al*. 2007) starting from leaves of 30-day-old plants grown at 22°C/18°C (day/night cycle) with 60% relative humidity and a photoperiod of 8/16 h (240 μmol m^−2^ s^−1^ photon flux density) in a combination of type-T soil (F.-E. Typ Nullerde, Hawita) and sand with a ratio of 3:1. Protoplasts were enzymatically released from leaves in a solution containing 0.4 M mannitol, 20 mM KCl, 20 mM MES, pH 5.7, 1% (w/v) cellulase R10, and 0.25% (w/v) macerozyme R10 (Duchefa Biochemie B.V.). Before use, the enzyme solution was heated to 55°C for 10 min to solubilize the enzymes. Subsequently, 10 mM CaCl_2_ and 0.2% BSA were added before the solution was filter sterilized with a 45 µm filter (Merck SA). Ten to 15 leaves with the upper epidermis peeled off were shaken (50-60 rpm) in the activated enzyme solution for 1 h in the dark. Subsequently, protoplasts were collected by filtration through a nylon mesh and centrifuged at 100 x g at 4°C. The final concentration of mesophyll protoplasts used in each experiment was 500,000 ml^−1^.

### Design of circRNAs

The structure models of all circRNAs and putative target mRNAs were predicted using mfold (version 3.6, mfold_util 4.7 and RNA Folding Form V2.3; www.unafold.org; Waugh et al. 2002; Zuker 2003) and RNAfold (RNAfold web server, university of Vienna; http://rna.tbi.univie.ac.at; (Gruber et al. 2008; Lorenz et al. 2011; Mathews et al. 2004). The 30 nt antisense target sequence for the *GFP* gene in the 50 nt circRNA_GFP_ was retrieved from the *GFP*-*ORF* region (Fig. S1A). *GFP* mRNA accessibility was assessed with the software RNAup (Vienna RNA Package, http://rna.tbi.univie.ac.at). In addition, two non-specific circRNAs (with no *GFP* sequences) were produced, which contain a randomized 25 nt or 46 nt sequence with a common 20 nt backbone, forming a 45 nt circRNA_CTR1_ and 66 nt circRNA_CTR2_, respectively.

### Production of circRNAs and linRNAs

The syntheses of circRNAs were performed as described (Nielsen et al. 2022; Pfafenrot et al. 2021). Briefly, the antisense and the unspecific control sequences were inserted between two spacers consisting of three unrelated nucleotides between the constant backbone, and this arrangement assured the stem-loop formation in both the antisense and control sequences. The oligonucleotide sequences for the circRNA synthesis, including the T7 promoter sequence, were commercially synthesized (Sigma-Aldrich). circRNAs were produced by *in-vitro* transcription from annealed DNA oligonucleotide templates (Table S2) using HighScribe T7 high-yield RNA synthesis kit (NEB) along with ATP, CTP, UTP, GTP (7.5 mM each), GMP (30 mM, Merck), and RNaseOut (Thermo Fisher Scientific) at 37°C for 2 h. Before circularization, the template DNA was digested with RQ1DNase (Promega) at 37°C for 30 min. Transcripts were purified with a Monarch RNA purification kit (NEB) and quantified with a Qubit™ RNA broad-range assay kit (Thermo Fisher Scientific). The RNA transcript was circularized overnight at 16°C with 200 U of T4 RNA ligase (Thermo Fisher Scientific) in 200 µL T4 RNA ligase buffer containing 0.1 mg mL^−1^ BSA and RNaseOut (Thermo Fisher Scientific). Following this reaction, circRNA was cleaned by phenol/chloroform extraction (Roth) and ethanol precipitation.

Single-stranded linear RNA (linRNA_GFP_) with the same nt sequence as circRNA_GFP_ was produced in the same way, but without circularization. Both linRNA and circRNA were further gel-purified from denaturing polyacrylamide gels as described (Breuer and Rossbach 2020). To confirm the circularity of circRNA, 250 ng of circRNA or linRNA were treated with or without 2 U of RNase R enzyme for 25 min at 37°C (Biozym) and analyzed by denaturing polyacrylamide gel electrophoresis followed by ethidium bromide staining.

### pGY1-35S::GFP:RFP plasmid construction

The red fluorescent protein (RFP) cassette was digested from the pBeaconRFP_GR vector (https://gatewayvectors.vib.be/index.php/ID:3_20, Bargmann and Birnbaum 2009) using the restriction enzyme *Nde*I (NEB, R0111S) for 20 min at 37°C followed by a deactivation step at 65° C for 10 min. pGY1-35S::GFP vector containing an *Nde*I restriction site was similarly digested, including a dephosphorylation with the enzyme Fast alkaline phosphatase (Thermo Fisher Scientific, EF0651). Both digestion products were run in a 1% agarose gel. Corresponding bands were excised from the gel, purified using Wizard^®^ SV Gel and PCR Clean-Up System (Promega), and ligated overnight at room temperature using T4 DNA ligase (Thermo Fischer Scientific, EL0011), resulting in the pGY1-35S::GFP:RFP vector.

### Transfection of protoplasts

Twenty µg of plasmid pGY1-35S::GFP:RFP (Fig. S2A) or pGY1-35S::GFP (Fig. S2B; Schweizer et al. 1999) along with circRNA or the respective linRNA were carefully added in a volume of 20 µl to 10^5^ mesophyll protoplasts in 200 µl MMg buffer (0.4 M mannitol, 15 mM MgCl_2_, 4 mM MES, pH 5.7). Then 220 µl of PEG-Ca^2+^ (40% PEG-4000, 0.2 M mannitol, 0.1 M CaCl_2_) were slowly added to the protoplast suspension and incubated at room temperature (Yoo *et al*., 2007). After 15 min of incubation, protoplasts were washed by centrifugation two times with W5 buffer (154 mM NaCl, 125 mM CaCl_2_, 5 mM KCl, 2 mM MES, pH 5.7). Subsequently, protoplasts (5× 10^5^ ml^−1^) were resuspended in W1 buffer (0.5 M mannitol, 20 mM KCl, 4 mM MES, pH 5.7) and incubated in the dark at room temperature. After 18 h, protoplasts were inspected under the fluorescence microscope (MZ16F Leica, Germany). At least three pictures were taken from the same treatment at different spots. Fluorescence was measured from these pictures by using ImageJ 1.54p software (https://imagej.net/ij/) and the ratio of fluorescence levels between the GFP fluorescent protoplasts (λ_exc_ 470, λ_em_ 525 nm) and total protoplasts (red auto fluorescence from the chlorophyll, λ_exc_ 480, λ_em_ 510 nm), or alternatively, RFP (λ_exc_ 550, λ_em_ 650 nm) was calculated.

### Plant material and growth conditions for callose deposition assay

*A. thaliana* Col-0 plants were grown from seeds in soil (LAT-Terra Standard Topferde Struktur 1b, Hawita, Vechta, Germany) complemented with 2,5 g/L fertilizer (Osmocote 12-7-19 +TE) and kept in a growth chamber equipped with LED lights under 12h/12h light/dark periods at 22°C/18°C. Leaf disks were excised using a cork borer and incubated overnight in 1 ml of water in the same chamber where the plants were grown. Leaf disks were then washed two times with water, placed on microscope slides, and covered with coverslips fixed with tape. The leaf disks were treated with 200 µl of a 0.1% aniline blue solution (pH=9) containing either water, poly(I:C) (Sigma-Aldrich) as a positive control, or different concentrations of circRNA or linRNA by adding the respective solution to the space between the slide and the coverslip and by evacuation for 2 min (0.08 MPa). After incubation in the dark for 30 min the callose staining at epidermal plasmodesmata was imaged with a Zeiss LSM 780 confocal laser scanning microscope equipped with ZEN 2.3 software (Carl Zeiss, Jean, Germany) by applying a 405 nm diode laser for excitation and filtering the emission at 475-525 nm. Eight-bit images acquired with a 40× 1.3 N.A. Plan Neofluar objectives with oil immersion were analyzed with ImageJ 1.53 software (https://imagej.net/ij/) using the plug-in calloseQuant (Huang et al. 2022). The fluorescence intensity levels of the callose spots were measured in 3-4 images taken from each leaf disk. Three leaf disks from three different plants were analyzed per condition. Normal distribution of the data was estimated and differences in p-values between treatments and the control (water) were determined by parametric one-way ANOVA followed by Dunnett’s multiple comparisons test using the Prism 8.4.0 software.

### Protein isolation, immunoblotting, and imaging

Protein extraction from protoplasts was performed using 4x SDS buffer (1 M Tris HCl, pH 6.8, 80% glycerol (v:v), bromophenol blue 10 mg, 4% SDS (w:v), 1 M of dithiothreitol (DTT) in 20 ml Milli-Q water). The sample was vortexed, heated to 95°C for 5 min and then centrifuged at 12,500 rpm for 2 min at 4°C. Protein concentration in the supernatant was determined according to the Bradford Ultra method (Bradford, 1976) on a Bio-Spectrophotometer (Eppendorf) at 595 nm. Then, 4 µg of each protein sample was loaded onto a 12.5 % SDS-PAGE gel. Afterwards, the proteins in the gel were transferred into the PVDF membrane (Merck KGaA) and blocked for one hour. Subsequently, the membrane was cut into two separate pieces based on the sizes of GFP (∼26.9 kDa) and Actin (∼45 kDa). The membrane containing the GFP band was incubated with the living colors monoclonal antibody JL-8, (Takara Bio Inc) and anti-mouse IgG-peroxidase conjugate (Sigma) as the secondary antibody. The membrane carrying the Actin band was incubated with an Actin polyclonal antibody (AS132640, Agrisera, Sweden) and goat anti-rabbit IgG HRP (AS09602, Agrisera, Sweden) as secondary antibody. After antibody incubation, the built-in software from the ChemiDoc MP imaging system (Bio-Rad) was used to evaluate protein band intensity in all western blots. Band intensity of control plants was set to one, and protein accumulation was calculated as a ratio between GFP and Actin. Protoplast counting for all the samples was done by ImageJ analysis.

### RNA isolation and gene expression analysis

Total RNA was extracted with the Direct-zol™ RNA Microprep kit (Zymo Research) and treated with DNase I following the manufacturer’s instructions. One µg or 500 ng of RNA was used for cDNA synthesis using a cDNA kit (RevertAid RT Kit, Thermo Fischer Scientific). GFP transcript levels were quantified by qPCR using SYBR Green JumpStart Taq ReadyMix (Sigma Aldrich, 1003444642) with a QuantStudio5 Real-Time PCR System (Applied Biosystems). The total volume of 10 µl and three technical replicates are considered for each reaction and 2 µl of ROX (CRX reference dye, Promega, C5411) was added to 1 ml of SybrGreen as a passive reference dye that allows fluorescent normalization for qPCR data. PCR conditions were 95°C for 5 min, followed by 40 cycles of 95°C for 15 s, 60°C for 30 s, and 72°C for 30 s, and then by a melting curve analysis. GFP expression levels were first normalized to RFP expression to account for transformation efficiency. Subsequently, fold changes of GFP expression were calculated using the ΔΔCt method (Livak and Schmittgen 2001) relative to the geometric mean of two endogenous housekeeping genes (Vandesompele et al. 2002), *Ubiquitin* (*UBQ5*, *AT3G62250*) and *Elongation Factor-1 alpha* (*EF1*α, *AT5G60390*), with protoplasts transformed with pGY1-35S::GFP:RFP vector serving as the control condition. One transformation of 100.000 protoplasts was considered as one biological replicate. The results of four biological replicates are included in the data analysis. The primer pairs employed for expression analysis are listed in Table S2.

### Analysis of ROS production

Leaf disks were collected from 4-weeks-old *N. benthamiana* plants (3 biological replicates) and incubated overnight in autoclaved Milli-Q water at 22°C in the dark. The following day, the water was replaced, and the incubation continued for additional 4 h. Leaf disks were transferred to a 96-well plate with 150 µL of a solution containing 17 µg/mL of luminol (Sigma-Aldrich) and 10 µg/mL horseradish peroxidase (HRP; Sigma-Aldrich) together with either 1 µM flg22 (ProteoGenix), 500 ng/µL of poly(I:C) (∼1 µM) (Sigma-Aldrich) or 17 ng/µL (∼1 µM) of linRNA_CTR2_ or circRNA_CTR2_. For the negative control, the elicitor was replaced by water. Luminescence detection was achieved using the microplate reader Varioskan LUX (Thermo Fisher Scientific) at 2 min intervals during 35 min. Mean values obtained from nine leaf disks per treatment were expressed as mean relative light units (RLU).

### MAPK phosphorylation of leaf extracts

The experiment was performed as described in Huang et al. (2023) with minor modifications. Arabidopsis leaf disks were collected from three plants and incubated overnight in autoclaved Milli-Q water at 22°C in the dark. After acclimatation, leaf disks were washed two times, then gently transferred to 96-well plate and incubated for an additional hour in autoclaved Milli-Q water. The water was replaced by 250 µL of a solution containing either 1 µM flg22 (EZBiolabs), or 50 ng/µL (∼3 µM) of linRNA_CTR1_ or circRNA_CTR1_. For the negative control, the elicitor was replaced by water. Leaf disks were vacuum infiltrated (0.08 MPa) for 30 min and immediately placed on liquid nitrogen. For the immunoblots, the frozen tissue was disrupted and resuspended in 100 µL 2X Laemmli buffer following a 5 min incubation at 95°C. Samples were separated in a 12% polyacrylamide gel and immunoblots were probed with antibodies against phosphor-p44/42 ERK (Cell Signaling Technology) and HRP-labeled secondary antibody (Thermo Fisher Scientific) for luminescence detection. Ponceau red was used for loading control.

### Statistical analysis

For Statistical analysis we used GraphPad Prism 8. The statistical significance between sets of parametric data was analyzed with either one sample Student’s *t*-test, or one-way ANOVA followed by Dunnett’s post-hoc test, whereas the one sample Wilcoxon test and Kruskal-Wallis tests were used for non-parametric sets. A description of the specific tests is given in figure legends. Callose quantification experiments were repeated twice with similar outcomes. All the other experiments were repeated at least three times.

## Acknowledgment

We thank Deutsche Forschungsgemeinschaft (DFG) and Agence National de la Research (ANR) for strong support. In later stages, this work was also supported by a grant from the Cercle Gutenberg (Alsace, France) to KHK and MH; and the Ernst-Leopold Klipstein Foundation, Paderborn Gießen, Germany, to SN.

## Supplementary data and figure legends

**Figure S1** Topology of designer circRNAs and their target sequences in the *GFP* gene. (A) *GFP*-ORF region nucleotide sequence with color code: the 30 nt target sequence of the 50 nt *GFP* antisense circRNA_GFP_ is highlighted in red. Highlighted in blue is the CaMV35S promotor sequence; highlighted in yellow is the 5’UTR; red font represents the circRNA target sequence (30 nt); start codon ATG underlined in green; stop codon TAA underlined in red; 3’UTR highlighted in green; terminator sequence highlighted in grey. (B) The exact position of the target sequence was selected based on the secondary structure model of the ORF. The red box inside the *GFP* mRNA structure indicates the binding site for the designated antisense circRNA_GFP_. (C) The selection of this region was confirmed by measuring mRNA accessibility using the RNAup software (Vienna RNA Package, http://rna.tbi.univie.ac.at/). The *GFP* mRNA sequence (5’UTR and 3’UTR included) was used with its own antisense RNA sequence to determine RNA accessibility. This analysis served as a guide for selecting sequence sections for antisense circRNA design. Green box indicates higher mRNA accessibility, chosen for the target sequence of the antisense circRNA_GFP_. (D) Sequences and structures of three circRNAs used in this study. The secondary structures of designer circRNAs were determined with the program mfold (www.unafold.org; Zuker 2003). The *GFP* target sequence is colored in green, while randomized control sequences are colored in blue. Respective nt sequences of all circRNA are shown in Table S1.

**Figure S2** Map of plasmids for Arabidopsis protoplast transfection. (A) pGY1-35S::GFP:RFP: The plasmid was prepared by using the pGY1-35S::GFP (Schweizer et al. 1999; see B) plasmid backbone, where RFP was inserted by cloning. (B) pGY1-35S::GFP backbone: The whole construct confers resistance to ampicillin and carbenicillin antibiotics. *GFP* is inserted and its expression is under the control of *Cauliflower Mosaic Virus* 35S (CaMV35S) promoter and terminator. The Maps were generated by SnapGene freeware (https://en.freedownloadmanager.org/Windows-PC/SnapGene-Viewer-FREE.html.

**Figure S3** Microscopic imaging of GFP fluorescence in Arabidopsis protoplasts. Protoplasts were transfected with 20 µg of plasmid pGY1-35S::GFP and 4 µg of *GFP* antisense circRNA_GFP_, or 4 µg of non-targeting circRNA_CTR1_. (A). After 18 hpt, protoplasts were examined under the microscope using two distinct filters to analyze the ratio of fluorescence levels between the GFP fluorescent protoplasts (λexc 470, λem 525 nm) and red chlorophyll autofluorescence (λ_exc_ 480, λ_em_ 510 nm). Fluorescence intensity was quantified based on images by using ImageJ 1.54p software. (B). The ratio between green pixels (GFP fluorescence) and red pixels (chlorophyll fluorescence) as calculated with ImageJ represented in the graph. The bar represents the measurements of ≥6 individual pictures taken at various positions. Statistical analysis was performed using one-way ANOVA, where ‘*’ denotes p ≤ 0.05 significance to the protoplast transfected without circRNA (control).

**Figure S4** Microscopic imaging of the dose dependency of GFP abundance in Arabidopsis protoplasts in response to circRNA treatment. Protoplasts were transfected with 20 µg of plasmid pGY1-35S::GFP and the indicated amount of circRNA_GFP_ (A) or circRNA_CTR1_ (B), respectively. At 18 hpt, protoplasts were inspected under the fluorescence microscope using two different filters to calculate the ratio in fluorescence levels between the GFP fluorescent protoplasts (λ_exc_ 470, λ_em_ 525 nm) and total protoplasts (red chlorophyll autofluorescence, λ_exc_ 480, λ_em_ 510 nm). Fluorescence was measured based on pictures by using ImageJ 1.54p software.

**Figure S5** Microscopic imaging of the GFP fluorescence in Arabidopsis RNAi mutants. Protoplats of mutants *dcl1-11* (A) and *ago1-27* (B) were transfected with 20 µg of plasmid pGY1-35S::GFP and 4 µg of *GFP* antisense circRNA_GFP_ or non-targeting circRNA_CTR1_. At 18 hpt, protoplasts were inspected under the fluorescence microscope using two different filters to calculate the ratio in fluorescence levels between the GFP fluorescent protoplasts (λ_exc_ 470, λ_em_ 525 nm) and total protoplasts (red chlorophyll autofluorescence, λ_exc_480, λ_em_ 510 nm). Fluorescence was measured based on pictures using ImageJ 1.54p software.

**Figure S6** Immunoblot analysis of the GFP abundance in Arabidopsis protoplasts of RNAi mutants *dcl2,3,4*, *ago2-1* and *ago4-1* upon treatment with circRNA_GFP_ or non-targeting circRNA_CTR1_. Protoplasts were transfected with 20 µg of pGY1-35S::GFP plasmid and 4 µg circRNA_GFP_ or circRNA_CTR1_, respectively. Equal amounts of protein were analyzed by Ponceau-S staining.

**Figure S7** RT-qPCR analysis of the amount of *GFP* transcripts in Arabidopsis protoplasts from RNAi mutants upon treatment with circRNA. Protoplasts were transfected with 20 µg of pGY1-35S::GFP plasmid and 4 µg of circRNA_GFP_ or non-targeting circRNA_CTR1,_ respectively. Relative *GFP* expression was measured after 18 hpt. Values were normalized to the housekeeping gene *Ubiquitin*. Bars represent an average of three independent biological experiments pooled together with standard error of the mean (SEM). No statistically significant differences between treatments and genotypes were detected (one sample *t*-test for (A), (B), (C), and (E) and one sample Wilcoxon test for (D), p≥ 0.05).

**Figure S8** Mean values of callose intensity levels at PD in individual images taken of epidermal cells of leaf disks treated with a 0.1% aniline blue solution containing water (control), 500 ng/µL of poly(I:C), or either 50 ng/µL or 250 ng/µL of circRNA_CTR2_. Error bars show the standard error of the mean. Parametric mean value variances were tested by one-way ANOVA followed by Dunnett’s multiple comparisons test. ***, p< 0.001; ns = non-significant.

**Table S1** Nucleotide sequence of circRNAs used in this study.

**Table S2** List of oligonucleotides used in this study.

Oligonucleotides are used for circRNA and dsRNA production, as well as for RT-qPCR.

## Statements & Declarations

### Funding

This work was funded by the Deutsche Forschungsgemeinschaft (DFG) in the research unit 5116 (exRNA) to KHK and AB/PS. It was also funded by ERA-NET SusCrop 2 program (DFG grant 459501999 to KHK and Agence National de la Research (ANR) grant ANR-21-SUSC-0003-01 to MH) as part of the project BioProtect coordinated by MH and carried out under the second call of the ERA-NET Cofund SusCrop, being part of the Joint Programming Initiative on Agriculture, Food Security and Climate Change (FACCE-JPI). SusCrop has received funding from the European Union’s Horizon 2020 research and innovation programme under grant agreement No 771134. In later stages, this work was also supported by a grant from the Cercle Gutenberg (Alsace, France) to KHK and MH. SN was partly supported by the Ernst-Leopold Klipstein Foundation, Paderborn Gießen, Germany.

### Competing Interests

The authors have no relevant financial or non-financial interests to disclose.

### Author Contributions

All authors contributed to the study conception and design. Material preparation, data collection and analysis were performed by M. Hossain, C. Pfafenrot, S. Nasfi, A. Sede, J. Imani, E. Šečić, A. Bindereif A, M. Heinlein, M. Ladera-Carmona and KH Kogel. The first draft of the manuscript was written by KH Kogel and all authors commented on previous versions of the manuscript. All authors read and approved the final manuscript.

### Data Availability

The datasets generated during and/or analysed during the current study are available from the corresponding author on reasonable request.

## Notes

### Competing Interest Statement

The authors have declared no competing interest.

### Summary of Updates

A new Fig. 6 and S8 shows that circRNA has no immunogenic activity.

