## Supplementary material for "Designer antisense circRNA_GFP_ reduces GFP abundance in Arabidopsis protoplasts in a sequence-specific manner, independent of RNAi pathways": Tables S1 and S2

**Table S1:** Nucleotide sequences of circRNAs and linRNAs used in this study.

| Name | Sequence (5’-3’) |
| --- | --- |
| ^1^circRNA_GFP_ | GGGAGUAAGCUCGUGCUGCUUCAUGUGGUCGGGGUAGCGGGCUUACAGUA |
| circRNA_CTR1_ | GGGAGUAAGCAGAUGCGCACCGCACAGAUGCGCACGCUUACAGUA |
| circRNA_CTR2_ | GGGAGUAAGCAAAAGUCAGUGAGUCAGUGUAAUACGGGAGGAUACCCGCUGUCAAAGCUUACAGUA |

^1^ This sequence is identical to the linRNA_GFP_ sequence. Underscored nt are needed for circularization.

**Table S2:** List of oligonucleotides used in this study.

| Primers Name | Sequences of  primers/oligonucleotides | Application |
| --- | --- | --- |
| *GFP*_F | GACGTAAACGGCCACAAGTTC | qPCR |
| *GFP*_R | AAGTCGTGCTGCTTCATGTG | qPCR |
| *At* Ubiquitin_F | GCTTGGAGTCCTGCTTGGACG | qPCR |
| *At* Ubiquitin_R | CGCAGTTAAGAG GACTGTCCGGC | qPCR |
| *AtEF1-α_F* | CTGTTGTAACAAGATGGATGCC | qPCR |
| *AtEF1-α_R* | CCCTCGAATCCAGAGATTGG | qPCR |
| RFP_F | GCGAGATCAAGATGAGGCTGA | qPCR |
| RFP_R | TAGTCCTCGTTGTGGGAGGT | qPCR |
| Antisense_circRNA_GFP__F | TAATACGACTCACTATAGGGAGTAAGCTCGTGCTGCTTCATGTGGTCGGGGTAGCGGGCTTACAGTA | circRNA production |
| Antisense_circRNA_GFP__R | TACTGTAAGCCCGCTACCCCGACCACATGAAGCAGCACGAGCTTACTCCCTATAGTGAGTCGTATTA | circRNA production |
| circRNA_CTR1__F | TAATACGACTCACTATAGGGAGTAAGCAGATGCGCACCGCACAGATGCGCACGCTTACAGTA | circRNA production |
| circRNA_CTR1__R | TACTGTAAGCGTGCGCATCTGTGCGGTGCGCATCTGCTTACTCCCTATAGTGAGTCGTATTA | circRNA production |
| circRNA_CTR2__F | TAATACGACTCACTATAGGGAGTAAGCAAAAGTCAGTGAGTCAGTGTAATACGGGAGGATACCCGCTGTCAAAGCTTACAGTA | circRNA production |
| circRNA_CTR2__R | TACTGTAAGCTTTGACAGCGGGTATCCTCCCGTATTACACTGACTCACTGACTTTTGCTTACTCCCTATAGTGAGTCGTATTA | circRNA production |
