## supplement figure for "Designer antisense circRNA_GFP_ reduces GFP abundance in Arabidopsis protoplasts in a sequence-specific manner, independent of RNAi pathways"

Fig. S1A

**>ORF_GFP_718_bp**

…TTTGGAGAGGACAGGGTACCCATCATACTAGTAGATCTGCGATCTAAGTAAGCTTGGCATTCCGGTACTGTTGGTAAAGCCACC**ATG**GTGAGCAAGGGCGAGGAGCTGTTCACCG
GGGTGGTGCCCATCCTGGTCGAGCTGGACGGCGACGTAAACGGCCACAAGTTCAGCGTGTCCGGCGAGGGCGAGGGCGATGCCACCTACGGCAAGCTGACCCTGAAGTTCATCTGC
ACCACCGGCAAGCTGCCCGTGCCCTGGCCCACCCTCGTGACCACCTTGACCTACGGCGTGCAGTGCTTCGC**CCGCTACCCCGACCACATGAAGCAGCACGA**CTTCTTCAAGTCCGC
CATGCCCGAAGGCTACGTCCAGGAGCGCACCATCTTCTTCAAGGACGACGGCAACTACAAGACCCGCGCCGAGGTGAAGTTCGAGGGCGACACCCTGGTGAACCGCATCGAGCTGA
AGGGCATCGACTTCAAGGAGGACGGCAACATCCTGGGGCACAAGCTGGAGTACAACTACAACAGCCACAAGGTCTATATCACCGCCGACAAGCAGAAGAACGGCATCAAGGTGAAC
TTCAAGACCCGCCACAACATCGAGGACGGCAGCGTGCAGCTCGCCGACCACTACCAGCAGAACACCCCCATCGGCGACGGCCCCGTGCTGCTGCCCGACAACCACTACCTGAGCAC
CCAGTCCGCCCTGAGCAAAGACCCCAACGAGAAGCGCGATCACATGGTCCTGCTGGAGTTCGTGACCGCCGCCGGGATCACTCTCGGCATGGACGAGCTGTACAAG**TAA**CTCGACT
AGAGTCGGGGCGGCCGGGGATCCTCTAGAGTCGACCTGCAGGCATGCCGCTGAAATCACCAGTCTCTC…

In principle, I am not sure if the journal allows different subpanels in separate sheets (all a matter of font size and presentation. I had never separated subpanels like this. Only if all subpanels are on a single page, you can be sure how it will look in the paper.

Fig. S1B

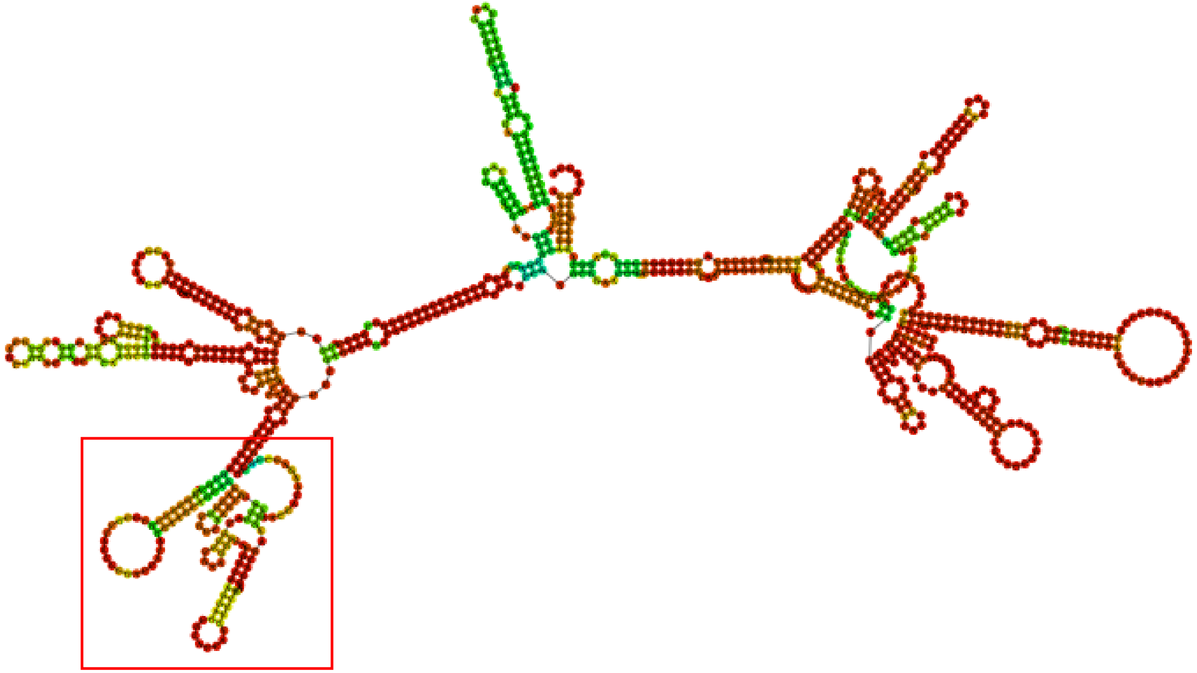

Fig. S1C

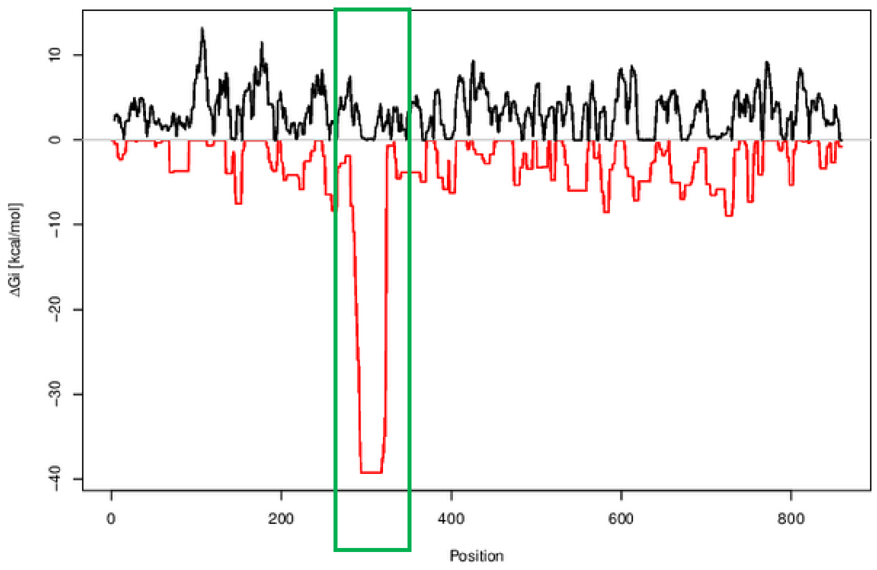

Fig. S1D

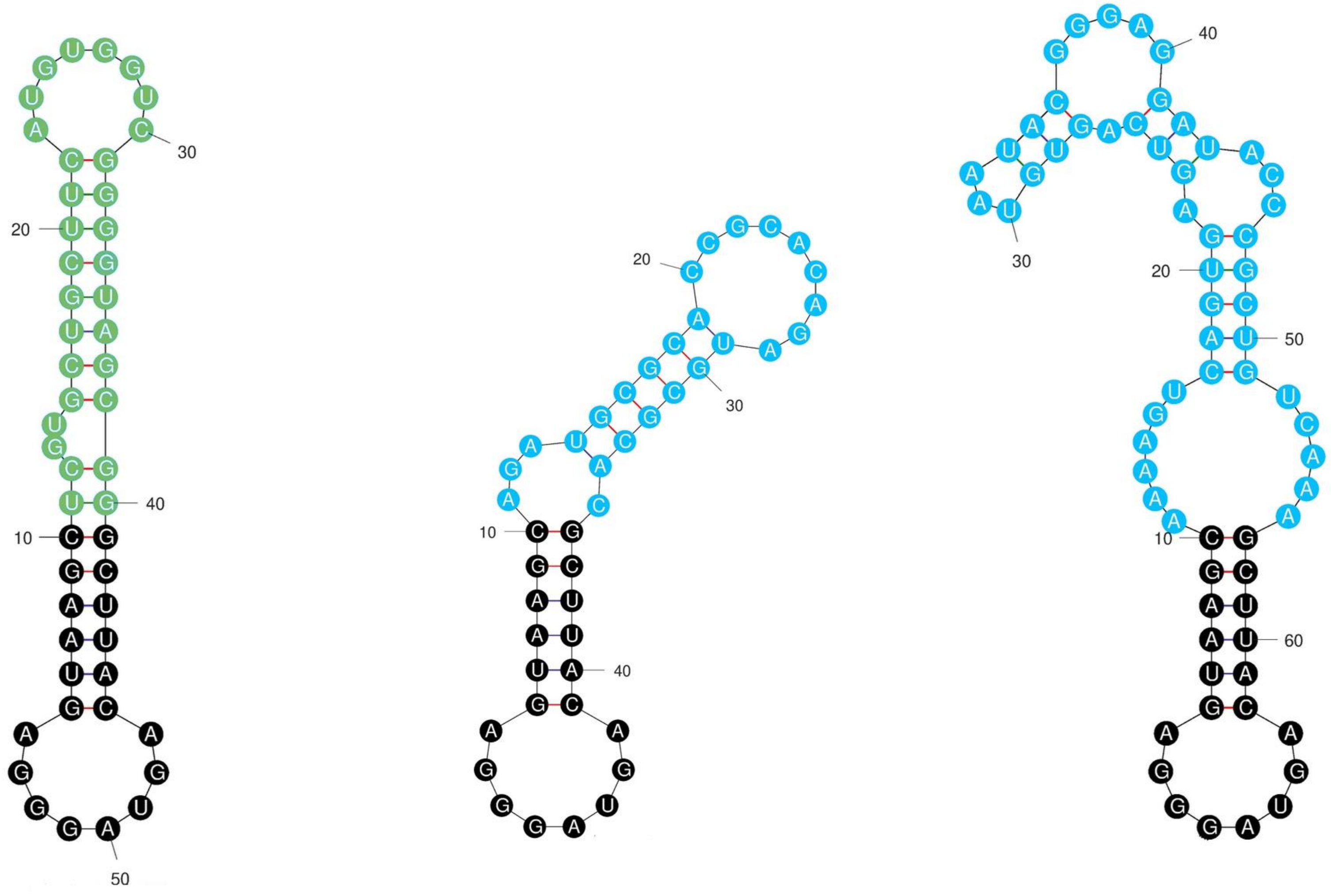


circRNA$_{GFP}$ circRNA$_{CTR1}$ circRNA$_{CTR2}$

Fig. S2A, B

A

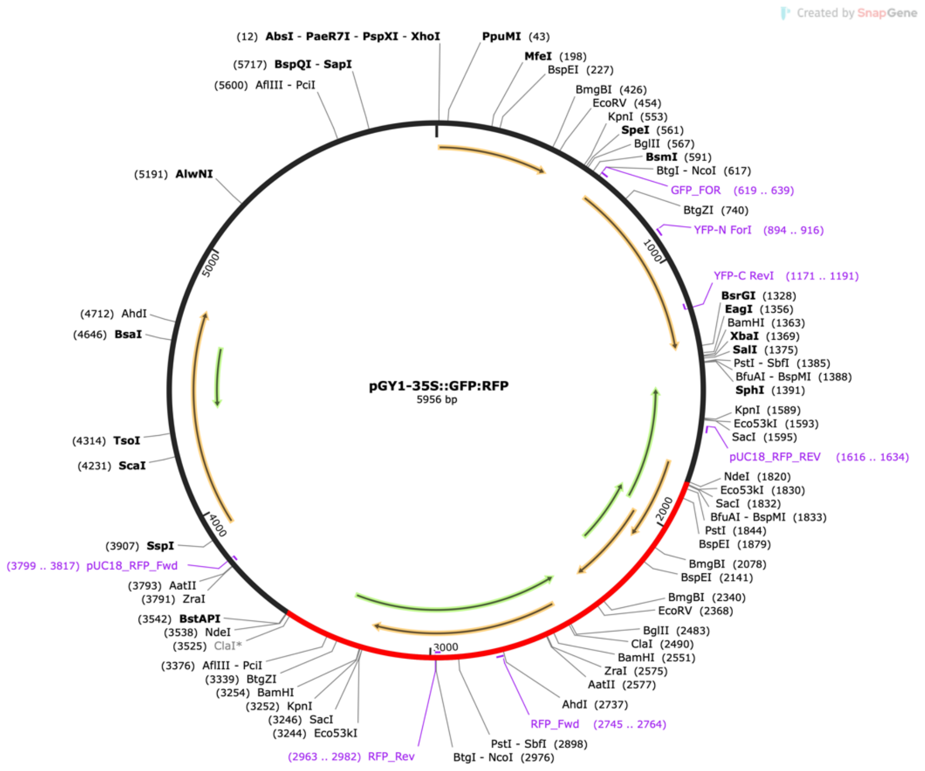


B

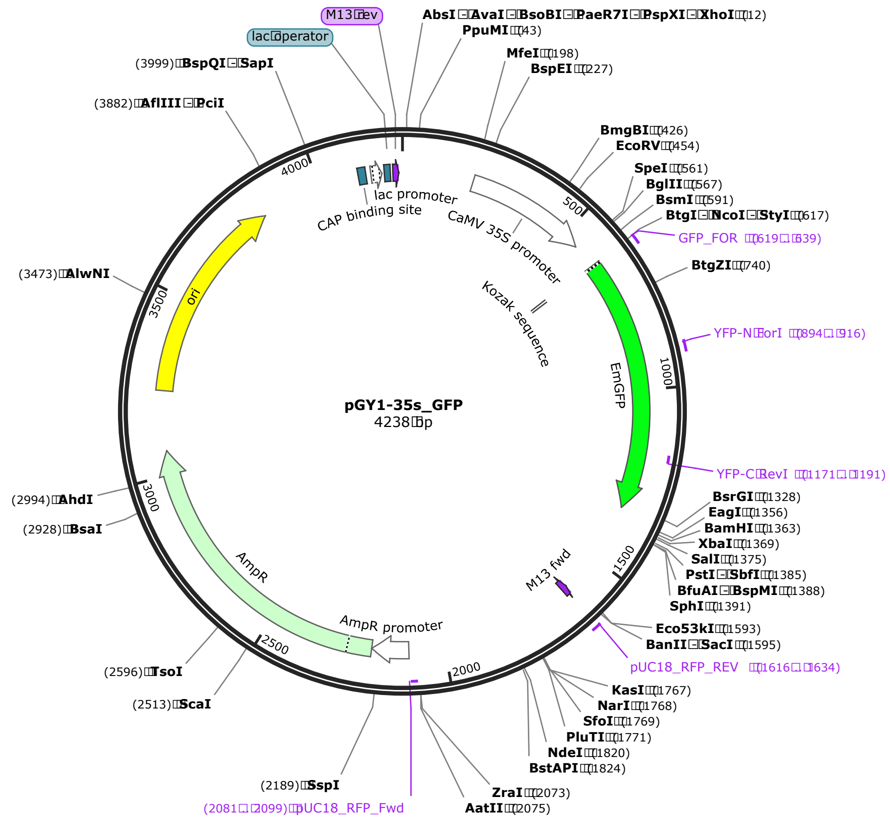

Fig. S3

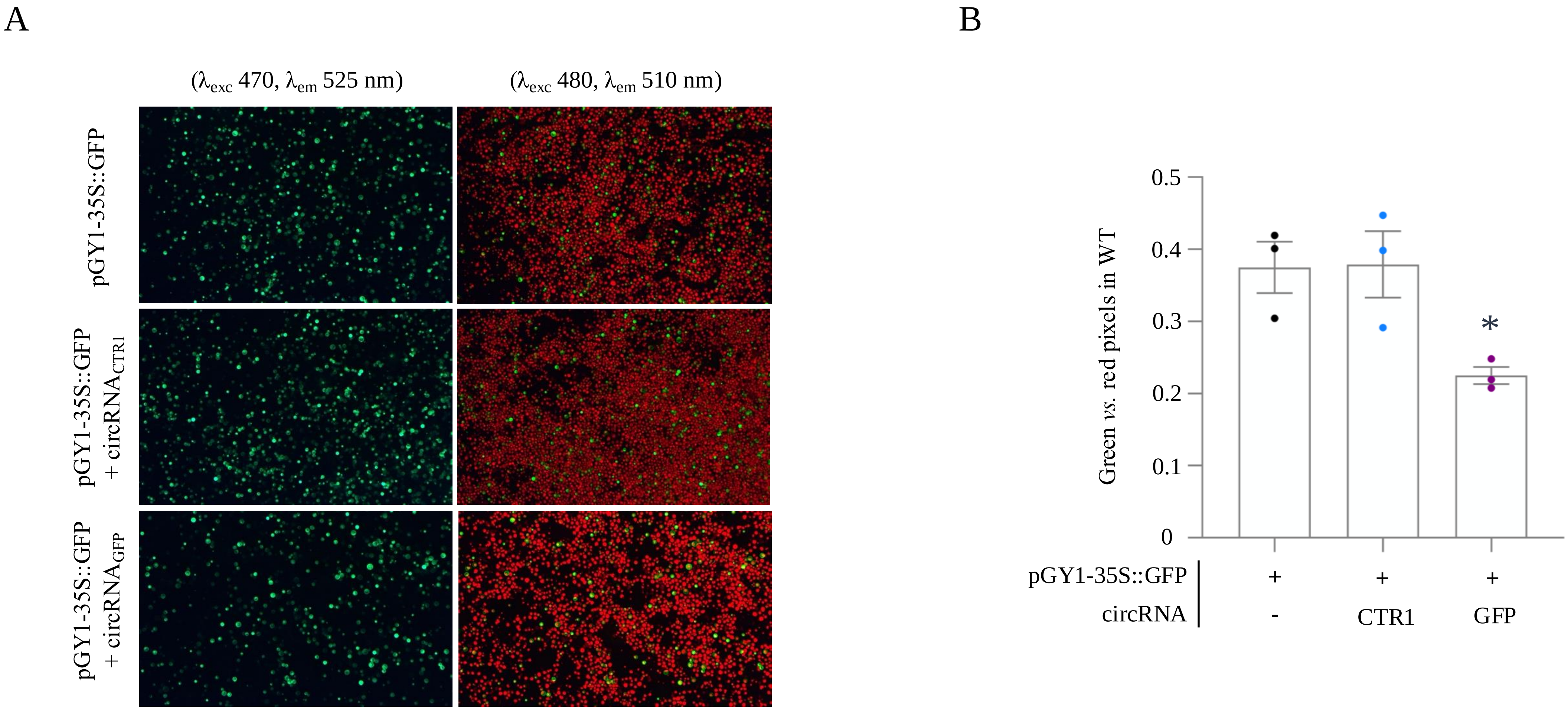
A
($\lambda_{exc}$ 470, $\lambda_{em}$ 525 nm)
($\lambda_{exc}$ 480, $\lambda_{em}$ 510 nm)
pGY1-35S::GFP
pGY1-35S::GFP + circRNA$_{CTR1}$
pGY1-35S::GFP + circRNA$_{GFP}$
B
Green *vs.* red pixels in WT
0.5
0.4
0.3
0.2
0.1
0
*
pGY1-35S::GFP
circRNA
+
+
+
-
CTR1
GFP

Fig. S4

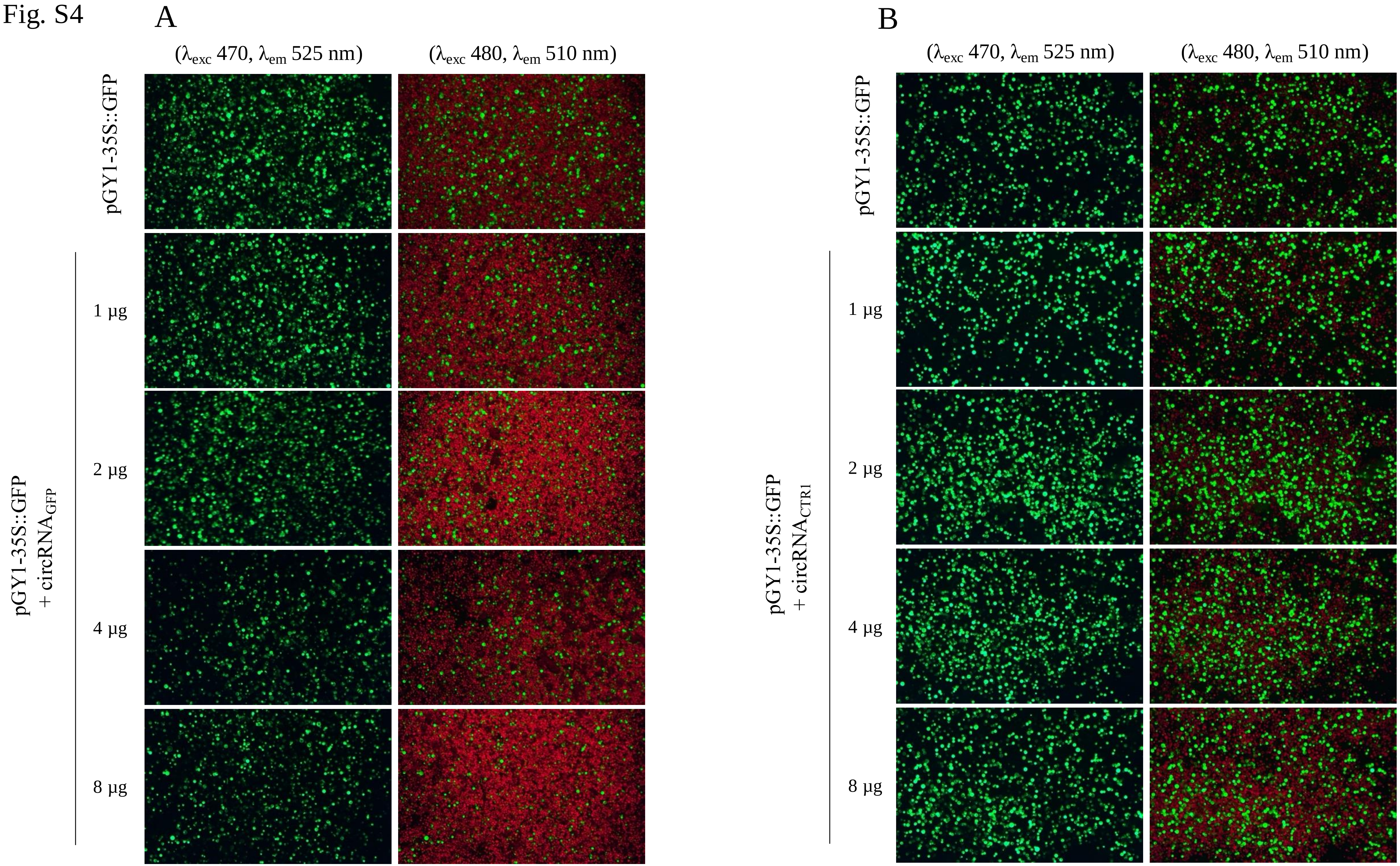
A
(λexc 470, λem 525 nm)
(λexc 480, λem 510 nm)
pGY1-35S::GFP
pGY1-35S::GFP
+ circRNAGFP
1 μg
2 μg
4 μg
8 μg
B
(λexc 470, λem 525 nm)
(λexc 480, λem 510 nm)
pGY1-35S::GFP
pGY1-35S::GFP
+ circRNACTR1
1 μg
2 μg
4 μg
8 μg

Fig. S5

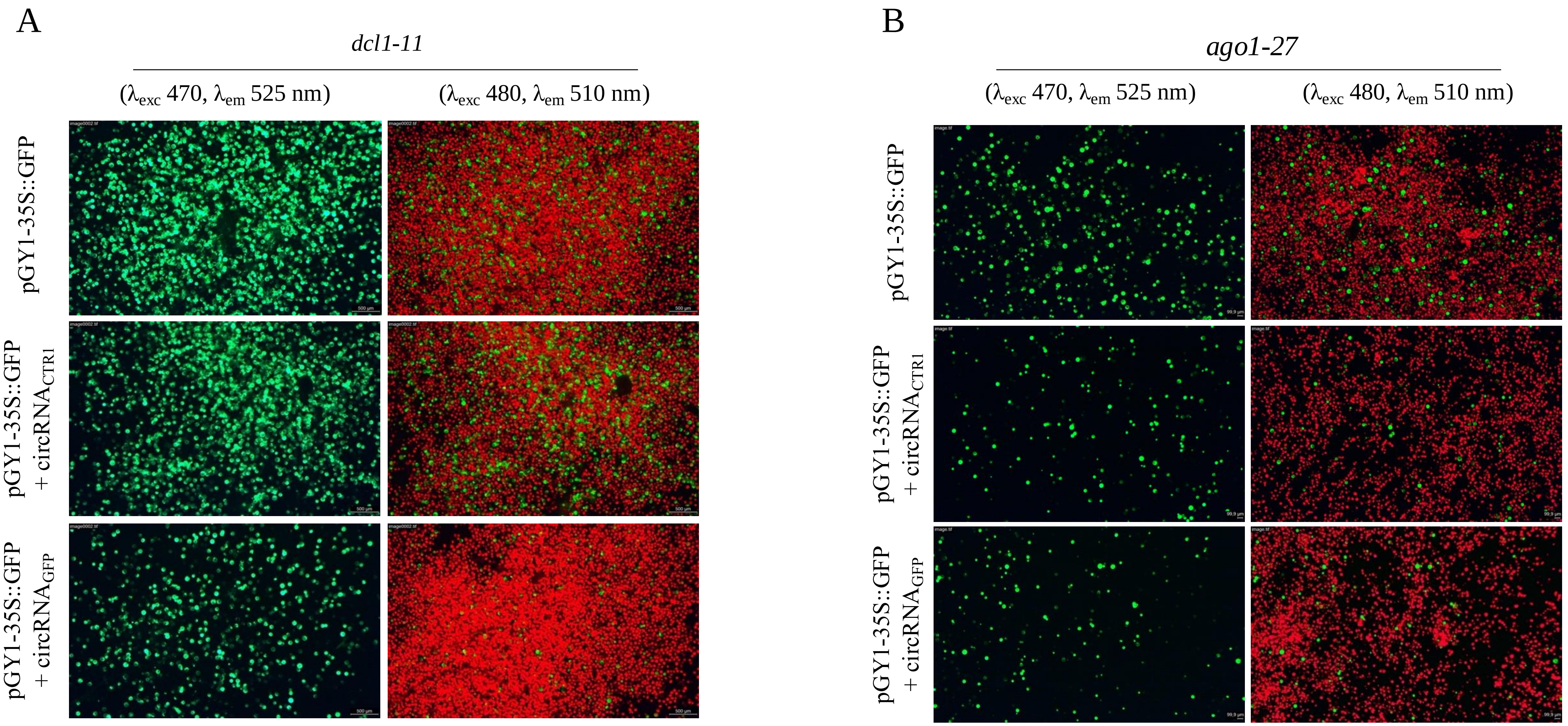
A
dcl1-11
(λexc 470, λem 525 nm)
(λexc 480, λem 510 nm)
pGY1-35S::GFP
pGY1-35S::GFP + circRNACTR1
pGY1-35S::GFP + circRNAGFP
B
ago1-27
(λexc 470, λem 525 nm)
(λexc 480, λem 510 nm)
pGY1-35S::GFP
pGY1-35S::GFP + circRNACTR1
pGY1-35S::GFP + circRNAGFP

Fig. S6

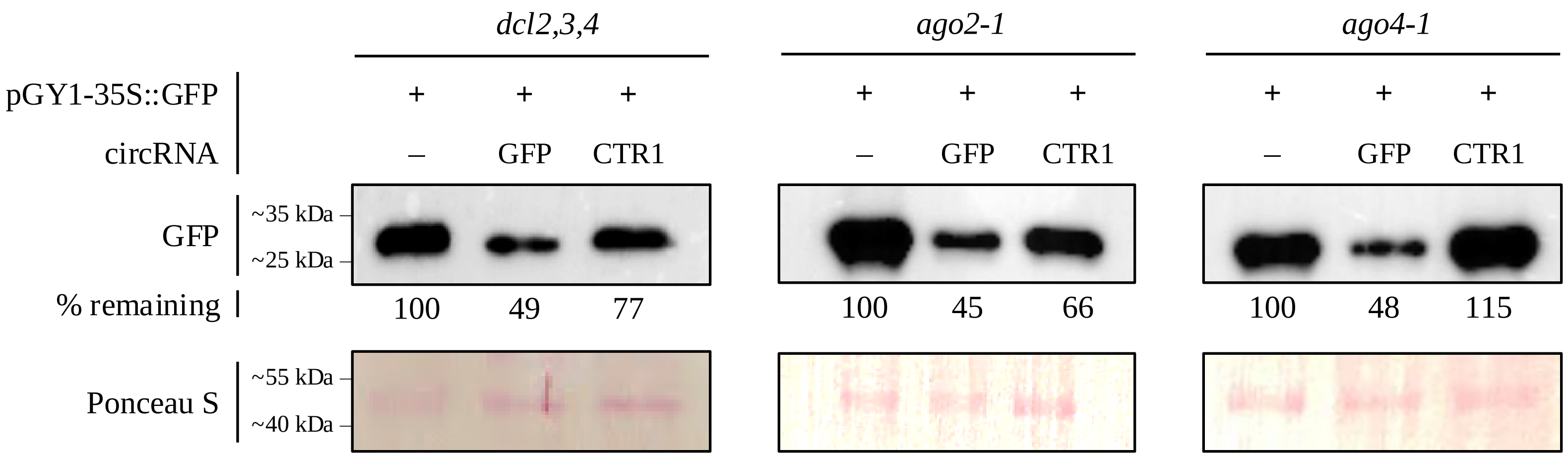

dcl2,3,4
ago2-1
ago4-1
pGY1-35S::GFP
+ + +
+ + +
+ + +
circRNA
– GFP CTR1
– GFP CTR1
– GFP CTR1
GFP
~35 kDa
~25 kDa
% remaining
100 49 77
100 45 66
100 48 115
Ponceau S
~55 kDa
~40 kDa

Fig. S7

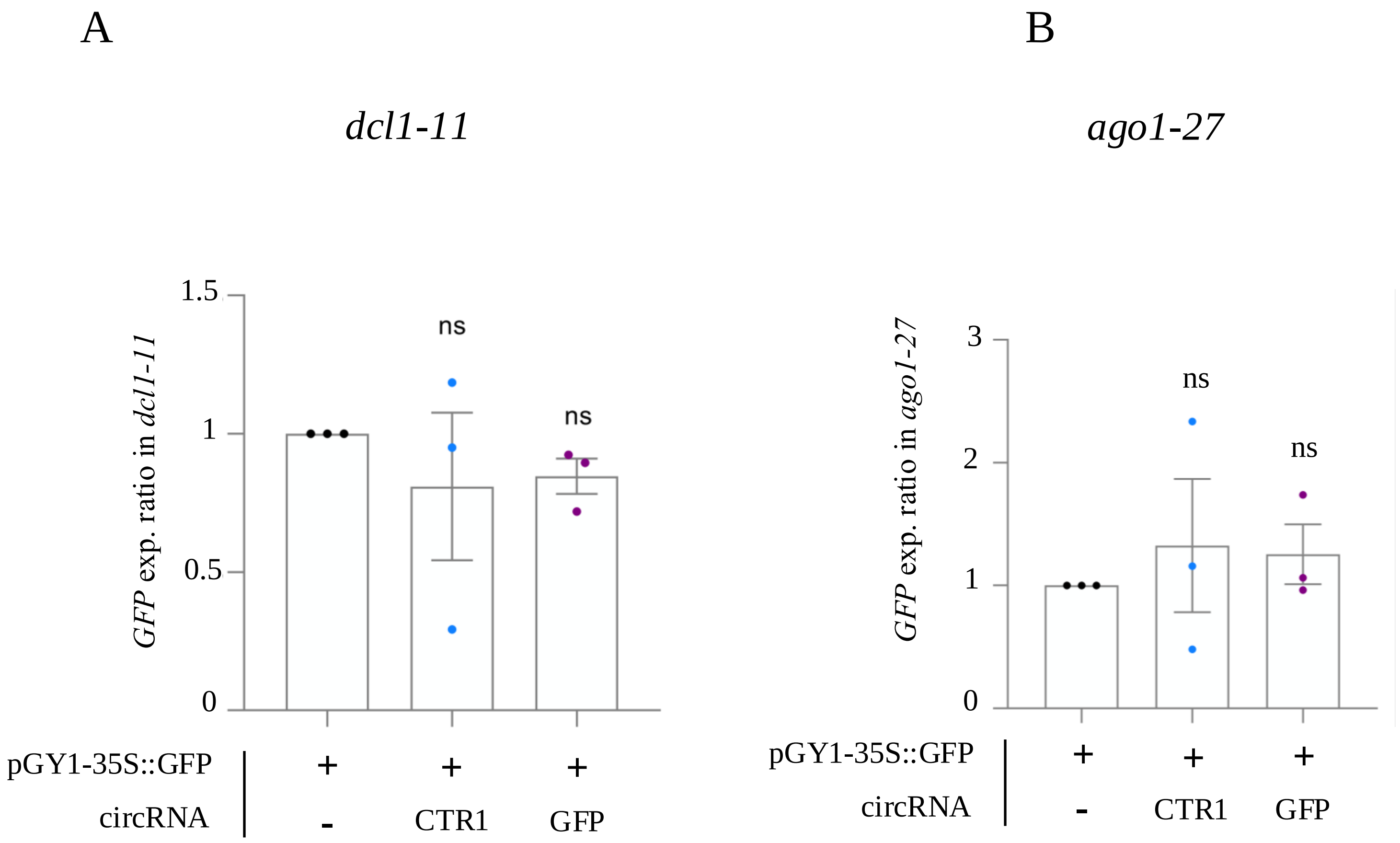
A
dcl1-11
GFP exp. ratio in dcl1-11
1.5
1
0.5
0
ns
ns
pGY1-35S::GFP
circRNA
+
+
+
-
CTR1
GFP
B
ago1-27
GFP exp. ratio in ago1-27
3
2
1
0
ns
ns
pGY1-35S::GFP
circRNA
+
+
+
-
CTR1
GFP

Fig. S7

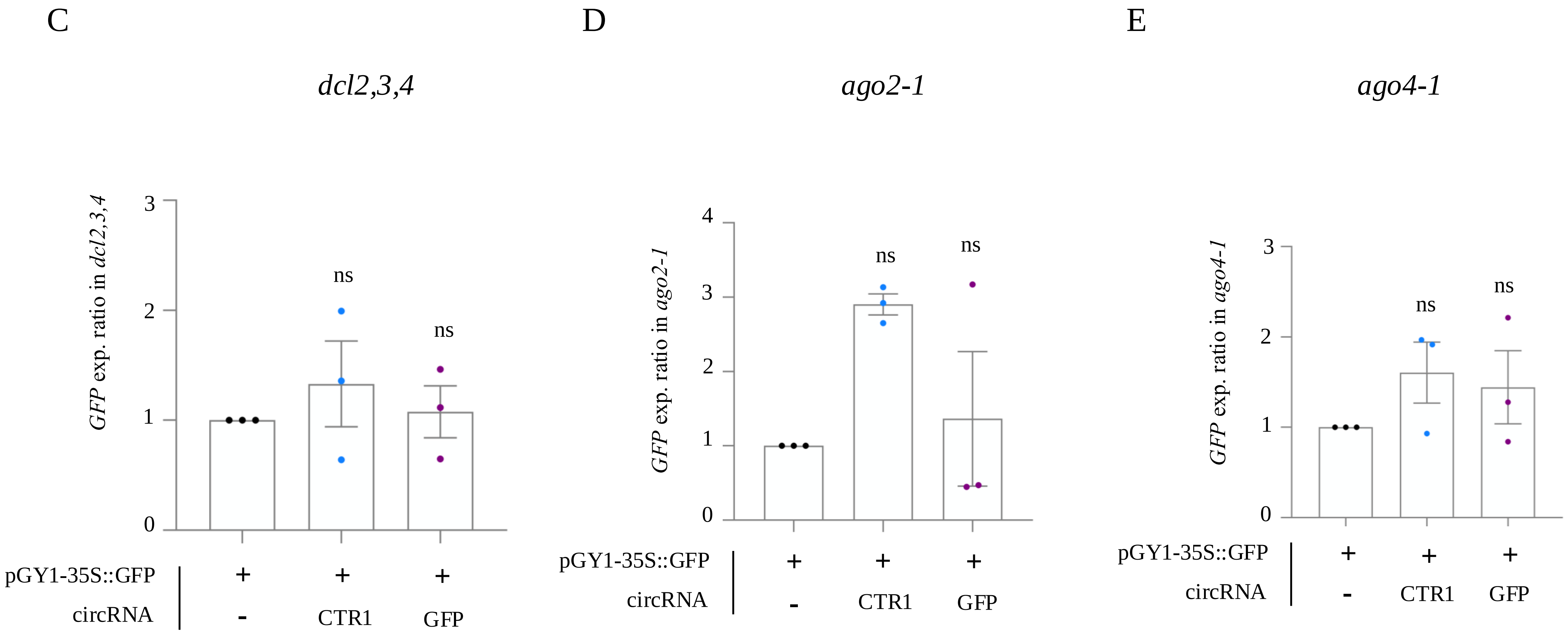
C
D
E
dcl2,3,4
ago2-1
ago4-1
GFP exp. ratio in dcl2,3,4
GFP exp. ratio in ago2-1
GFP exp. ratio in ago4-1
ns
pGY1-35S::GFP
circRNA
+
-
CTR1
GFP

Fig. S8

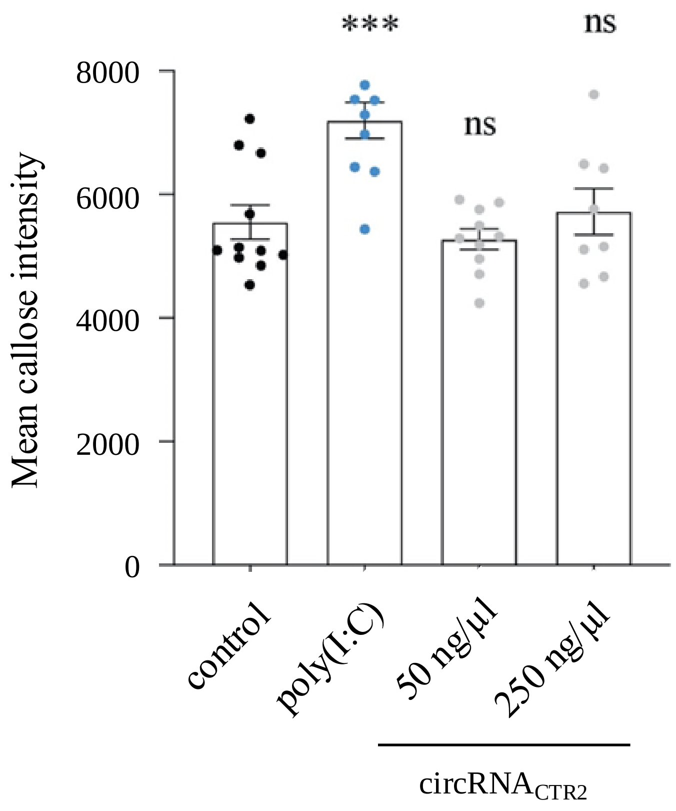

***
ns
ns
Mean callose intensity
8000
6000
4000
2000
0
control
poly(I:C)
50 ng/µl
250 ng/µl
circRNACTR2
